# Anti-capsule human monoclonal antibodies protect against hypervirulent and pandrug-resistant *Klebsiella pneumoniae*

**DOI:** 10.1101/2024.02.14.580141

**Authors:** Emanuele Roscioli, Vittoria Zucconi Galli Fonseca, Soraya Soledad Bosch, Ida Paciello, Giuseppe Maccari, Giampiero Batani, Samuele Stazzoni, Giulia Cardinali, Giusy Tiseo, Cesira Giordano, Yuwei Shen, Laura Capoccia, Matteo Ridelfi, Marco Troisi, Noemi Manganaro, Chiara Mugnaini, Dario Cardamone, Concetta De Santi, Annalisa Ciabattini, Linda Cerofolini, Marco Fragai, Danilo Licastro, David P. Nicolau, Kamilia Abdelraouf, Simona Barnini, Francesco Menichetti, Marco Falcone, Claudia Sala, Anna Kabanova, Rino Rappuoli

## Abstract

The silent pandemic caused by antimicrobial resistance (AMR) requires innovative therapeutic approaches. Human monoclonal antibodies (mAbs), which are among the most transformative, safe and effective drugs in oncology and autoimmunity, are rarely used for infectious diseases and not yet used for AMR. Here we applied an antigen-agnostic strategy to isolate extremely potent human mAbs against *Klebsiella pneumoniae* (Kp) sequence type 147 (ST147), a hypervirulent and pandrug-resistant clonotype which is spreading globally. Isolated mAbs target the bacterial capsule and the O-antigen. Surprisingly, although both capsule- and O-antigen-specific mAbs displayed bactericidal activity in the picomolar range *in vitro*, only the capsule-specific mAbs were protective against fulminant ST147 bloodstream infection. Protection correlated with *in vitro* bacterial uptake by macrophages and enchained bacterial growth. Our study describes the only drug able to protect against pandrug-resistant Kp and provides a strategy to isolate mAbs and identify correlates of protection against AMR bacteria.

## INTRODUCTION

Antimicrobial resistance (AMR) has been enlisted by the World Health Organization, the European Medicines Agency and the United Nations as one of the top ten global health priorities, due to its impact on human health and socio-economic welfare worldwide^1^. Among AMR pathogens, *Enterobacteriaceae* causing threats in hospital settings are considered the most critical^2^ and are currently dominated by *Klebsiella pneumoniae* (Kp) species^3^. Most Kp strains that cause severe infections belong to two major pathotypes, namely the classical and hypervirulent ones^4^. Classical Kp is a frequent cause of healthcare-associated infections and can easily acquire mobile genetic elements associated with AMR^5^. In particular, multidrug-resistant Kp strains carrying extended-spectrum β-lactamases and carbapenemases have played a major role in global AMR spread. Among carbapenemases, the New Delhi Metallo-β-lactamases (NDM) are extremely alarming since they confer resistance to the most recent combinations of β-lactams and β-lactamase inhibitors (i.e., ceftazidime–avibactam, imipenem–relebactam, and meropenem–vaborbactam), which are considered the last line of defense^6^. NDMs are encoded by *bla_NDM_* genes located on large multi-resistance plasmids that have been spreading across high-risk AMR Kp clones throughout all continents^7,8^. On the other hand, hypervirulent Kp (hvKp) isolates carry large virulence plasmids encoding genes related to capsule synthesis and siderophore production^4^. Despite showing higher susceptibility to antimicrobial agents, hvKp strains can cause severe community-acquired infections, such as pyogenic liver abscess, endophthalmitis, and meningitis^7–10^, even in healthy individuals.

Although hypervirulence and multidrug resistance have evolved distinctly within Kp, convergence of both traits has been described recently, particularly in extensively drug resistant NDM-producing Kp strains^11^. Notably, an NDM-1-positive sequence type 147 (ST147_NDM-1_) Kp strain carrying both AMR and hypervirulence genes has been disseminating globally^12–14^ since mid-late 2000s and was associated with acquisition of carbapenemase-encoding genes in various countries^12–14^. The only antibiotic to which ST147_NDM-1_ is still sensitive is colistin which is associated with high systemic toxicity and is used as the last resort treatment for AMR infection^15–17^. Today, ST147 accounts for about 5% of Kp infections worldwide^18^. In 2018^19–21^ ST147 isolate landed in Tuscany (Italy), causing a nosocomial outbreak associated with a 30-day mortality rate close to 40%^19^. Genomic surveillance data indicate that ST147 isolates from the Tuscany outbreak carry a highly diversified plasmid content^19^, bearing numerous AMR genes and virulence factors. Similar genetic rearrangements had been already detected in Russia (2017)^22^, United Kingdom (2018-2019)^23^ Egypt (2019)^24^, and in the 2019 Kp outbreak in Germany^25^, where different Kp STs coharbouring virulence genes and *bla_NDM_* were reported. Indeed, an ancestor of the Tuscany outbreak strain^20^ has been recently classified as pandrug-resistant (PDR)^26,27^, leaving scarce therapeutic solutions against this pathogen.

Taking into consideration a scenario where high-risk Kp clones can cause catastrophic epidemics, it is mandatory to search for effective treatment strategies. In this context, human monoclonal antibodies (mAbs) represent a powerful tool that can rapidly progress to innovative prophylactic and therapeutic solutions. Notably, mAbs are unique in having intrinsically good safety profiles and avoiding impact on the host microbiota given their high specificity^28^. However, their use against the AMR threat has been underexplored. In fact, while 26 mAbs against bacterial pathogens have entered clinical phases for efficacy evaluation, only a few have gained FDA approval^28^, and none of them is directed against Kp.

At the preclinical level, several human and murine mAbs targeting Kp, but not ST147_NDM-1_ strains, have been reported. A few of these mAbs are directed against the capsular polysaccharide or the O-antigen and both types have shown protective efficacy against ST258 Kp in a murine intratracheal infection model^29,30^ or in models of acutely lethal pneumonia by ATCC8045 Kp^31^, bacteremia by ST258^32^ and O3a and O3b types of Kp (O3:K11)^33^. However, while the anti-capsule antibodies are clearly associated with protection in some studies^34^, the role of O-antigen antibodies is still controversial because other studies reported that the O-antigen may not be accessible to antibodies in capsulated strains^35,36^. Although valuable, the above findings find little applicability to ST147_NDM-1_ for one major reason: the astonishing variability of antigenic polysaccharidic structures across Kp strains. Indeed, more than 100 capsule serotypes (K antigen) have been described so far^37^, while 11 distinct O-antigen serogroups exist, with four O-antigens (O1, O2, O3, and O5) associated with 80-90% of Kp clinical isolates^4^. Moreover, continuous diversification of O-antigen structures, such as addition of pyruvate to O1b^38^ and GlcNAc to O2ac^39^, generates new epitopes. This imposes the development of serotype-specific mAbs or cross-validation of the pre-existing ones. On the other hand, mAbs targeting the type III fimbrial component MrkA were protective in a mouse model of lethal pneumonia caused by ATCC29011 Kp^40^ and could find broader application as MrkA is conserved across various Kp strains. However, another gap withholds their applicability against AMR Kp, specifically against the hypervirulent and pandrug-resistant ST147_NDM-1_: the lack of knowledge on the correlates of protection against bloodstream infection. Most mAb development efforts have focused on target-specific approaches, whereas no comprehensive studies comparing mAbs against larger panel of Kp epitopes have been undertaken.

In this work, we describe a powerful method for unbiased mAb selection against ST147_NDM-1_ not bound to a particular predetermined antigen, which allowed the isolation of potent bactericidal mAbs from donors that experienced ST147_NDM-1_ bloodstream infection. By using a combination of three high-throughput functional *in vitro* assays we defined functional properties of mAbs associated with protection against fulminant bloodstream infection. Our study describes the first fully human mAb against the hypervirulent and pandrug-resistant ST147_NDM-1_ Kp lineage and shows in a definitive manner that highly bactericidal anti O-antigen antibodies are not protective against this hypervirulent and pandrug-resistant strain. This is an example of how a rationally designed experimental pipeline allows to predict correlates of *in vivo* efficacy that could be universally applied to other emerging AMR strains.

## RESULTS

### Isolation of bactericidal human mAbs against Tuscany outbreak strain ST147_NDM-1_

To isolate Kp-specific mAbs representing the breadth of antibody repertoire induced by Kp infection, an antigen-agnostic approach was used. Peripheral blood mononuclear cells (PBMCs) were collected from 7 convalescent patients who recovered from ST147_NDM-1_ Kp bloodstream infection of the Tuscany nosocomial outbreak (**Table S1**). A total of 18,390 CD19^+^CD27^+^IgM^-^IgD^-^ memory B cells (MBCs), either IgG- or IgA-positive, was single-cell sorted and plated together with CD40L-expressing feeder cells, IL-2 and IL-21 to promote MBCs activation, proliferation and antibody secretion (**Figure 1A; Figure S1A**). MBCs culture supernatants were screened by enzyme-linked immunosorbent assay (ELISA) against ST147_NDM-1_ bacteria to select Kp-reactive mAbs. 214 mAbs specific for antigens displayed on the bacterial surface were identified (**Figure 1B)**.

**Figure 1.**
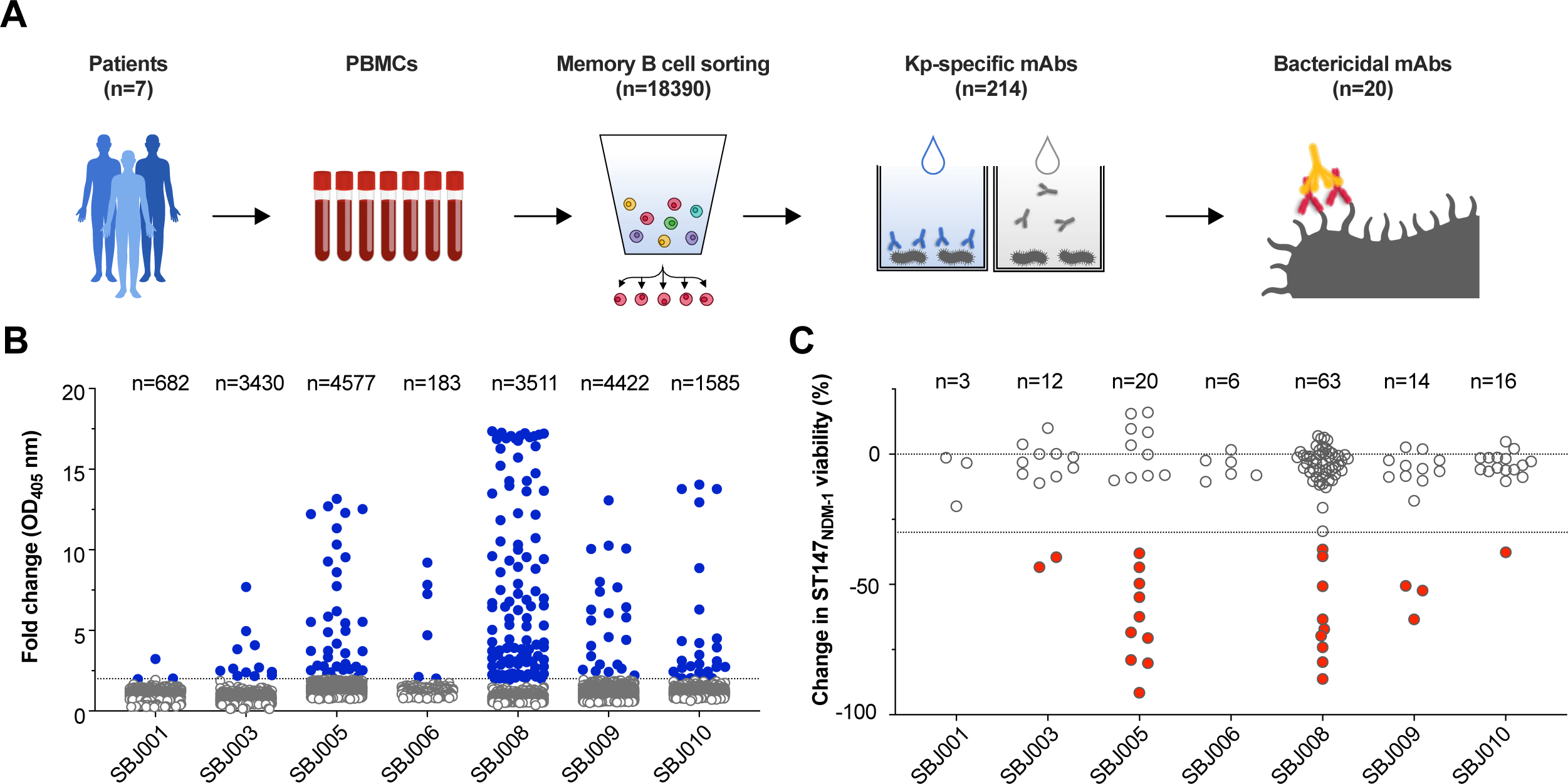
Isolation and selection of Kp-specific mAbs against ST147_NDM-1_ Tuscany outbreak strains. (A) Schematic workflow of anti-Kp antibody isolation and selection. (B) ELISA screening of supernatants of 18,390 single cell-sorted MBCs isolated from 7 convalescent patients. The assay was performed to detect IgG/IgA against surface antigens of ST147_NDM-1_ c.i.1 and ST147_NDM-1_ c.i.2. 214 MBCs supernatants showed a signal at least 2-fold above the blank of OD_405_ and were thus selected for further analysis (blue dots). (C) L-SBA on recombinant mAbs against ST147_NDM-1_ c.i.1. 30% reduction in bacterial viability was used as a cut-off for selection. 25 mAbs showing bactericidal properties were identified (red dots).

Upon molecular cloning of their V_H_ and V_L_ variable portions (**Table S2**), sequences of 134 Kp-reactive mAbs were recovered with successful V_H_ and V_L_ pairing. To validate binding properties of the isolated mAbs, they were expressed on a small scale and tested by ELISA against two outbreak-belonging ST147_NDM-1_ Kp isolates (**Figure S1B**). Further, mAbs were screened for their ability to induce complement-dependent killing of ST147_NDM-1_ Kp^41^. For this purpose, we developed a high-throughput luminescence-based serum bactericidal assay (L-SBA) that measures ATP content as a proxy of Kp viability^42,43^. Four dilutions per mAb were tested in the presence of an exogenous source of complement and antibodies causing a reduction greater than 30% in bacterial viability were considered as positive hits. Out of the 134 Kp-binding mAbs screened, 25 candidates showing bactericidal activity against ST147_NDM-1_ were selected (**Figure 1C**) and expressed on the larger scale. Upon excluding five mAbs that failed to re-confirm their bactericidal activity, binding and functional properties of the remaining 20 bactericidal ST147_NDM-1_-targeting mAbs were evaluated.

### Kp strain library profiling reveals that bactericidal mAbs display different degrees of cross-reactivity and antigen specificity

To characterize specificity and cross-reactivity of the isolated bactericidal mAbs, flow cytometry was employed to profile their binding against a panel of bacterial strains. Pathogenic Kp belonging to different genetically distant STs were included (**Table 1, Figure S2A**), as well as other *Klebsiella* species and several commensal strains. Antibody binding to bacterial surface displayed different levels of intensity, suggesting differential expression of target antigens or variable affinity of the mAbs (**Figure 2A**). Moreover, mAbs showed high specificity to pathogenic *sensu stricto* Kp strains as they did not react with other *Klebsiella* species nor with commensals (**Figure S2B**). Based on their binding profiles, mAbs were grouped into two main clusters. The first cluster (c*luster 1*) encompassed mAbs with rather restricted binding features, recognizing exclusively the outbreak strain ST147 carrying either *bla*_NDM-1_ or *bla*_NDM-9_ genes^44^. The second group of mAbs (c*luster 2*) displayed a broader binding pattern, being able to recognize up to seven different sequence types of pathogenic Kp (**Figure 2A**).

**Figure 2.**
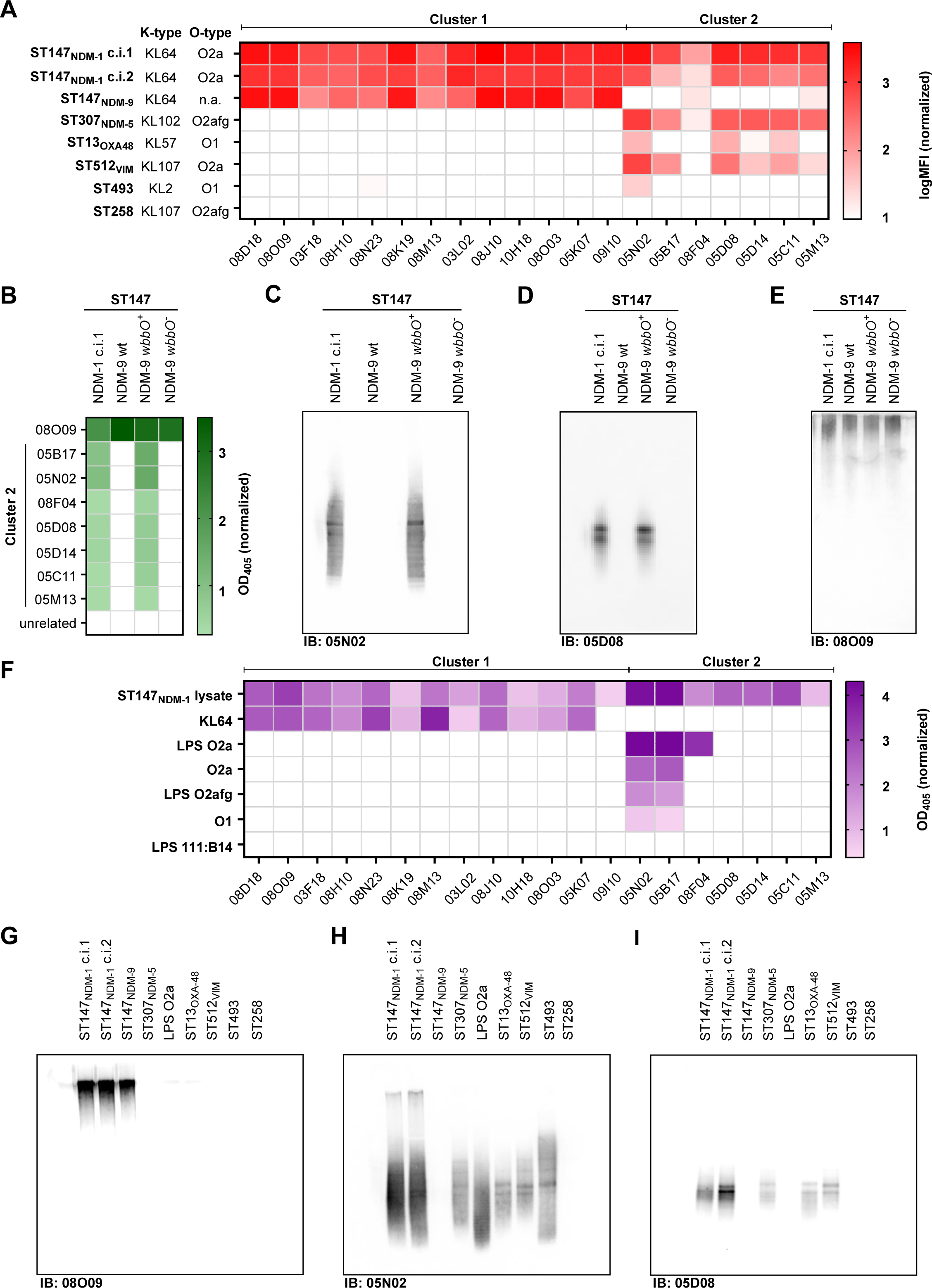
Binding properties of 20 bactericidal ST147_NDM-1_-specific mAbs. (A) Heatmap showing high-throughput flow cytometry screening of 20 functional mAbs against a panel of genetically diverse Kp strains. MFI was normalized to no mAb controls and transformed in logarithmic scale. (B) Heatmap of ELISA values obtained with 08O09 (cluster 1 mAb) and cluster 2 mAbs against ST147_NDM-1_ c.i.1, ST147_NDM-9_ (wt), ST147_NDM-9_/pSEVA23a1_*wbbO* (*wbbO^+^*), ST147_NDM-9_/pSEVA23a1 (*wbbO^-^*) strains. (C-E) Immunoblot (IB) of 05N02 (panel C), 05D08 (panel D) and 08O09 (panel E) against ST147_NDM-1_ c.i.1, ST147_NDM-9_ (wt), ST147_NDM-9_/pSEVA23a1_*wbbO* (*wbbO^+^*), ST147_NDM-9_/pSEVA23a1 (*wbbO^-^*) strains. (F) Heatmap of ELISA with purified capsule and O-antigens. Unrelated *E. coli* LPS O111:B4 was used as negative control and total bacterial lysate as a positive control. Values at least 3-fold above the blank of OD_405_ were considered as positive hits. (G-I) Representative immunoblots (IB) of selected anti-Kp mAbs probed against total sugar extracts of indicated strains. 08O09 (panel G) recognizes high molecular weight KL64 capsule. 05N02 (panel H) recognizes O-antigens with broad molecular weight distribution belonging to O2- and O1 types. 05D08 (panel I) recognizes medium molecular weight O-antigen structures from O2-carrying Kp strains.

**Table 1.**
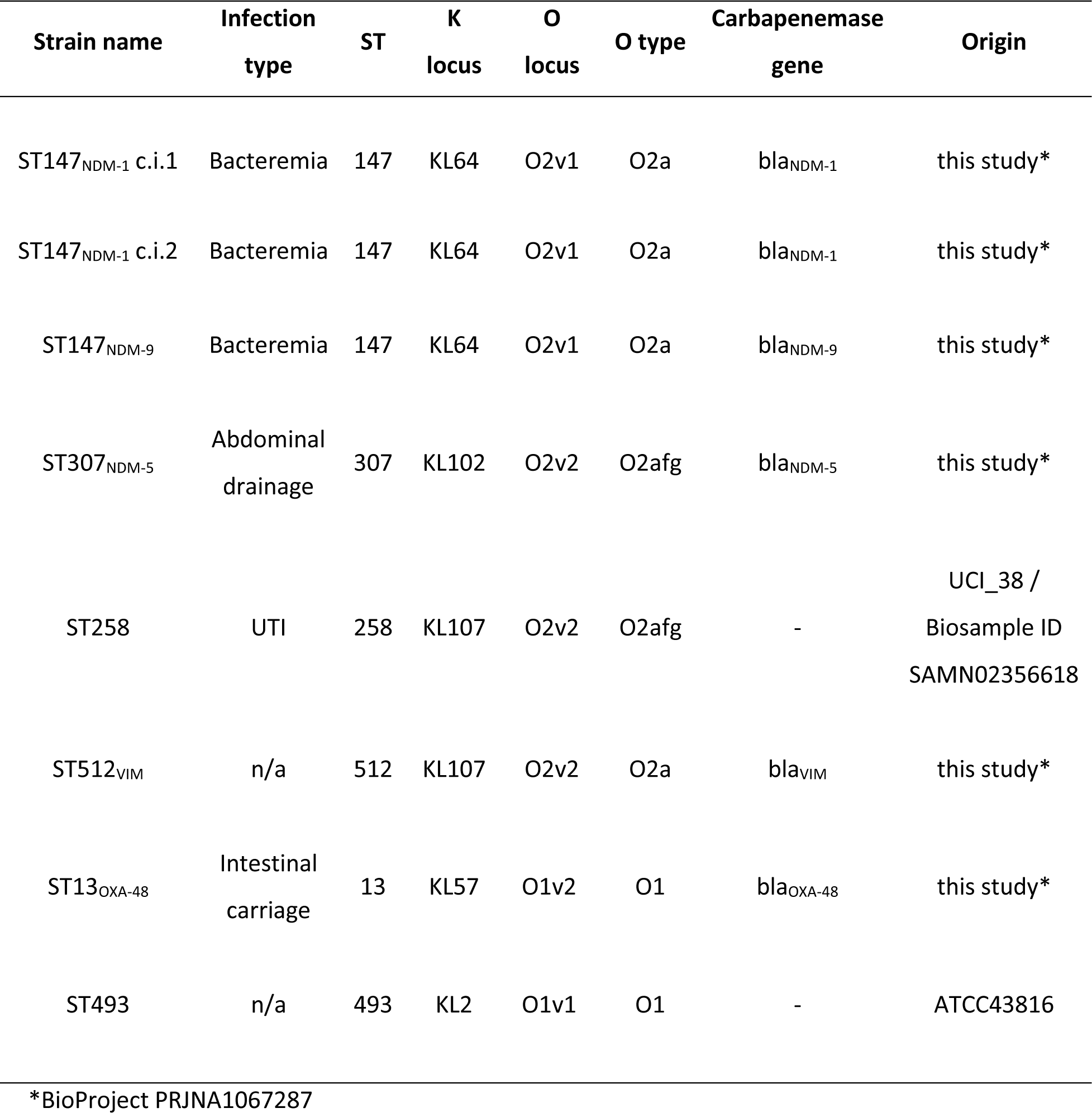
List of Kp isolates used in the manuscript. ST, K locus, O locus and presence of carbapenemase gene information were predicted from genomic analysis.

Comparative whole genome sequencing of the Kp strain library allowed inferring the surface antigens targeted by the selected mAbs. Pangenome analysis of the isolates identified a subset cluster of genes (COG) exclusively present in strains recognized by cluster 1 mAbs (**Table S3**). Notably, COG associated with the KL64 locus of *wcaJ* was identified. WcaJ functions as the initiating enzyme in the synthesis of colanic acid, a primary constituent of the capsule of *Enterobacteriaceae*^45^. This finding aligned with the specificity of cluster 1 mAbs, which exclusively recognized strains carrying genes belonging to the KL64 biosynthetic locus (**Figure 2A**)^46^, thus suggesting that their cognate antigen might be the capsule.

Regarding the putative target of cluster 2 mAbs, we observed that they reacted with Kp strains belogning to different STs but displayed almost null binding to ST147_NDM-9_ (**Figure 2A**). Therefore, we assumed that these mAbs were very likely directed against a surface antigen lacking in the ST147_NDM-9_ isolate. Genomic analysis of the O-antigen biosynthetic loci revealed that ST147_NDM-9_ presented a single-point deletion in position 474 of the *wbbO* gene (**Figure S3A**), which encodes for a glycosyltransferase essential for O2a biosynthesis (**Figure S3B**)^47^. This mutation causes frameshift and premature termination of translation likely hindering the production of the complete O-antigen^48^. Indeed, HPLC and SDS-PAGE analyses of total sugar extracts from ST147_NDM-9_ confirmed that this strain was a natural O-antigen-deficient mutant when compared to ST147_NDM-1_ (**Figure S4A and B**). Thus, we hypothesized that cluster 2 mAbs recognized the ST147_NDM-1_ O-antigen. To confirm this, ST147_NDM-9_ was transformed with a plasmid that carried the wild type *wbbO* gene, or with the empty vector as a control. ELISA experiments demonstrated that binding of cluster 2 mAbs was restored in the merodiploid strain ST147_NDM-9_ *wbbO*^+^, whilst background-level signals were measured ST147_NDM-9_ *wbbO*^-^ carrying empty vector (**Figure 2B**). Consistent with these results, immunoblotting analysis of total sugar extracts showed recognition of ST147_NDM-9_ *wbbO*^+^ by the cluster 2 mAbs 05N02 (**Figure 2C)** and 05D08 (**Figure 2D**), while no difference among strains was observed with the cluster 1 mAb 08O09 (**Figure 2E**). Overall, these data confirmed successful complementation of ST147_NDM-9_ and identification of the O-antigen as the target of cluster 2 mAbs.

### ST147-specific bactericidal mAbs target ST147_NDM-1_ capsule and O-antigen

Biochemical validation of the mAb cognate antigens was sought by ELISA against purified Kp capsule **(Figure S4C and D)** and O-antigens of various origins (**Figure S4E, F and G**). KL64 capsule was recognized only by cluster 1 mAbs (**Figure 2F**). Moreover, immunoblotting of Kp total sugar extract further confirmed that these antibodies decorated high molecular weight (MW) structures of KL64-bearing Kp strains, in agreement with electrophoretic profiles of polysaccharidic capsules^49^ (**Figure 2G**).

Instead, ELISA against purified O-antigens revealed a more heterogeneous picture within cluster 2 mAbs. Three mAbs bound to purified LPS/O2a O-antigen, with variable degree of cross-reactivity against O2afg and O1 O-antigens (**Figure 2F**). Their immunoblotting evidenced a ladder-like binding pattern, compatible with typical O-antigen immunoblotting profiles^48^ (**Figure 2H**). Four other mAbs from cluster 2 did not recognize purified LPS or O-antigens in ELISA (**Figure 2F**). However they displayed a similar ladder-like binding profile against total sugar extracts of Kp strains expressing O2a O-antigens (**Figure 2I**)^48^ and, as described in the previous paragraph, successfully bound the *wbbO*-complemented ST147_NDM-9_ strain (**Figure 2B and D**). Altogether, these analyses allowed grouping bactericidal ST147_NDM-1_-targeting antibodies into KL64-specific and O2a O-antigen-specific mAbs, wherein some of the latter were found to be cross-reactive against O2afg and O1 type O-antigen.

High-resolution and high-content imaging indicated that mAbs directed against capsule and O-antigen displayed different localization patterns (**Figure 3A**). For instance, the anti-capsule mAb 08O09 showed higher intensity binding levels compared to the anti-O-antigen mAbs 05D08 and 05N02 (**Figure 3B)**. In addition, while 08O09 signal occupied a median area of 8.3±3.4 µm^2^, 05D08 and 05N02 mAbs distributed over a smaller area of 4.9±1.6 µm^2^ and 4.8±1.5 µm^2^, respectively (**Figure 3C**, **Figure S5**). Co-staining experiments with 08O09-A488 and 05N02-A555 conjugated mAbs confirmed that 08O09 localized on the outermost layer of the bacteria (**Figure 3D**). Whereas, in three-color assays, 05N02-A555 co-localized with 05D08-A647 and distributed more internally than 08O09-A488 (**Figure 3E**), in agreement with differential specificity.

**Figure 3.**
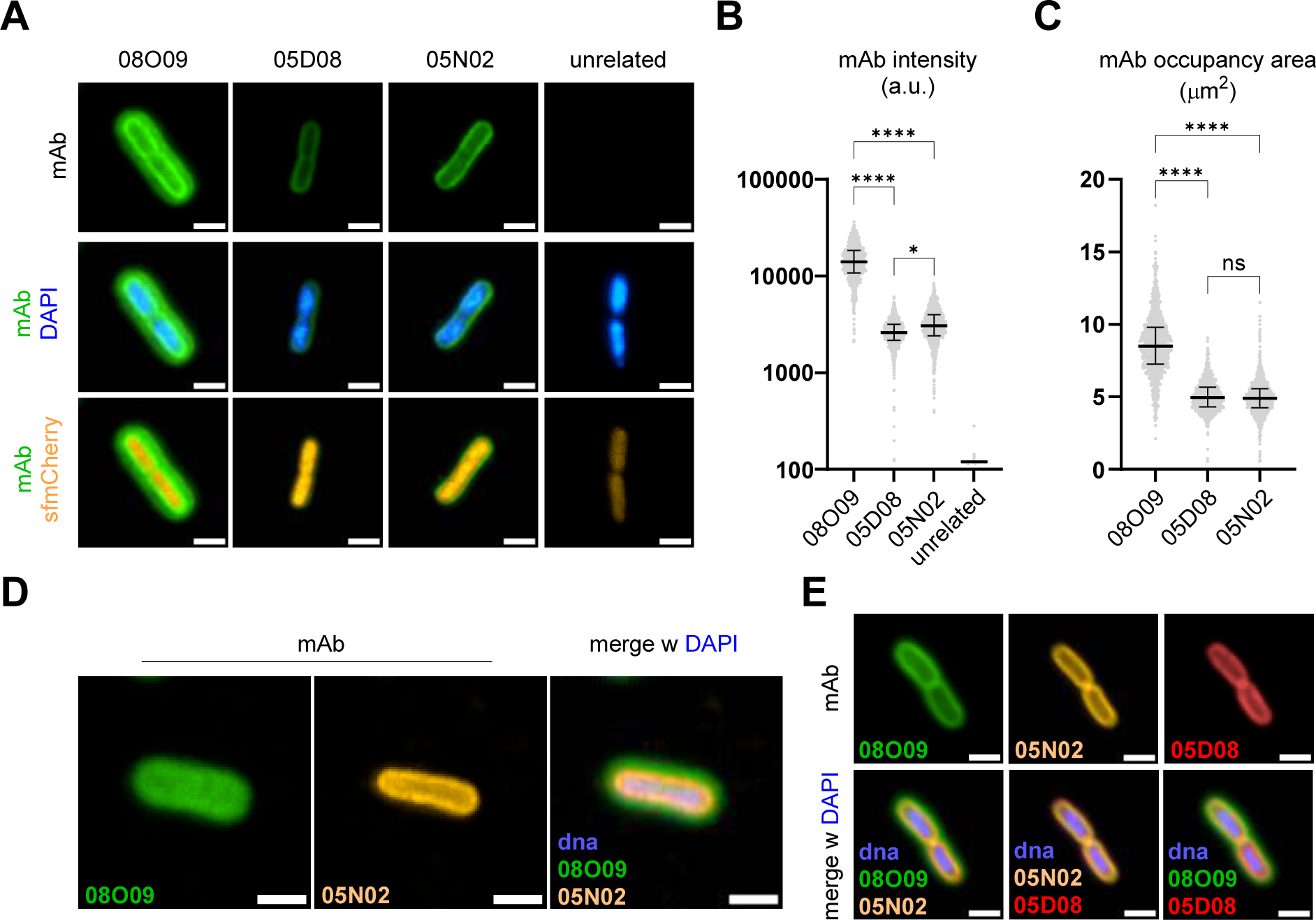
Microscopy characterization of anti-Kp mAb binding on ST147_NDM1_ c.i. 1 strain. (A) Images show ST147 _NDM-1_ c.i. 1 expressing sfmCherry stained with 08O09, 05D08 and 05N02 mAbs labelled with anti-human A488 conjugated secondary antibody (green). Bacterial DNA is stained with DAPI (blue). Scale bar 2 µm. (B) Scatter plot showing quartiles range of A488 intensity quantification at the single bacterium level (median intensity ± sd/number of bacteria are reported for each mAb: 12289±9278 a.u./3475 for 08O09; 2518±1250 a.u./3024 for 05D08; 2860±1601 a.u./3269 for 05N02; 119.1±3.7 a.u./3078 for unrelated mAb). Two-way ANOVA from 4 independent experiments, * p=0.0143; **** p<0.0001. (C) Scatter plot displaying quartiles range of the area of individual A488 spots (median area ± sd/number of spots are reported for each mAb: 8.3±3.4 µm^2^/3475 for 08O09; 4.9±1.6 µm^2^/3072 for 05D08; 4.8±1.5µm^2^/3310 for 05N02). Two-way ANOVA from 4 independent experiments, **** p<0.0001; ns, non-significant. (D) Images of ST147_NDM1_ c.i. 1 stained with 08O09-A488 (green), 05N02-A555 (yellow) conjugated mAbs and DAPI (blue). Scale bar 2 µm. (E) Representative images showing ST147_NDM1_ c.i. 1 stained with 08O09-A488 (green), 05N02-A555 (yellow) and 05D08-A647 (red) conjugated mAbs. DNA was stained with DAPI (blue). Scale bar 2 µm.

### mAbs sequence analysis reflects functional clustering

Analysis of heavy (HC) and light chain (LC) mAb sequences revealed insights into their germline composition. The genes encoding the HCs and LCs of 20 bactericidal mAbs were sequenced, and their IGHV and IGKV genes were analyzed (**Figure S6A**). The most common HC V gene was *IGHV3-23* (n=8, 40%), followed by *IGHV3-21* (n=3, 15%). The most prominent HC J gene was *IGHJ4* (n=15, 75%), mostly recombining with *IGHV3-23* (n=7, 35%), but showing diversity in the V gene. Most of the selected mAbs used the κ chain (n=17, 85%), with only three out of 20 using the λ chain. The light chain (LC) V gene usage was not as polarized as in the HC, with the *IGKV3-20* and the *IGKV2-30* used by 30% of the mAbs (n=3+3). Similarly, the LC J gene did not show polarized usage, with the most prominent V gene being *IGKJ1* (n=6; 30%), followed by *IGKJ2* (n=4, 20%). Initial examination evidenced a bias in germline usage for anti-capsular mAbs, *with IGHV3-23*04*;*IGHJ4*02* being the most common HC recombination (n=6, 30%) (**Figure S6B**), while two clonal pairs were detected among O-antigen-specific mAbs (*IGHV3-15*01*;*IGHJ4*02*, *IGHV1-18*01*;*IGHJ4*02*) (**Table S4**). To further investigate sequence relationships, the heavy and light chain sequences were analyzed separately using Clustal Omega, and sequence distances were calculated (**Figure S6C, D**). Within the HC alignment, two distinct clusters can be seen containing the anti-capsular mAbs using to the *IGHV3-23*04*;*IGHJ4*02* combination. Interestingly, these two groups contained other mAbs using the *IGHV3-23*04* or *IGHV3-21*01* V gene, suggesting an association of these two V genes with anti-capsular functionality. In the LC alignment there did not seem to be such a polarization, probably due to the more variegate LC recombination. Nonetheless, some functional preferences could be observed, with the LC V gene *IGKV1-17*01* and *IGKV1-27*01* being the most frequent anti-capsular combinations. The outcomes of these analyses agreed with previously described functional clustering, reinforcing the connection between sequence variations and functional attributes of the antibodies.

### Functional profiling reveals that isolated mAbs have extremely potent bactericidal activity, however only anti-capsular mAbs are poly-functional

To better assess protective properties of anti-capsule and anti-O-antigen bactericidal antibodies, several *in vitro* assays that could predict antibody-mediated protection *in vivo* were employed. Bactericidal mAbs were expressed as human IgG1 with the E430G mutation in the Fc domain, which enhances the hexamerization process that naturally occurs between Fc domains and thus potentiates antibody effector function^50^. The processivity of complement-dependent killing assays (SBA) was enhanced by optimizing a fluorescence-based SBA (F-SBA) in the 384-well format and by using resazurin metabolism as a readout of bacterial viability. F-SBA profiling of mAbs against complement-sensitive pathogenic Kp strains (ST147_NDM-1_, ST147_NDM-9_, and ST307_NDM-5_) revealed that most of the antibodies were functional in the picomolar range of concentrations, with the most potent mAb (05D08) showing about 0.5 ng/mL IC_50_ values (**Figure 4A; Figure S7A, B, C and D**). The ability to kill different Kp strains correlated with mAbs cross-binding properties showed in Figure 2A, since anti-KL64 mAbs (cluster 1) were effective against KL64-bearing ST147_NDM-1_ and ST147_NDM-9_, while anti-O-antigen mAbs (cluster 2) displayed activity against O2 O-antigen-bearing ST147_NDM-1_ and ST307_NDM-5_ (**Figure 4A**). F-SBA results allowed ranking mAbs according to their IC_50_ values and the extent of bacterial viability reduction (**Figure S7E**).

**Figure 4.**
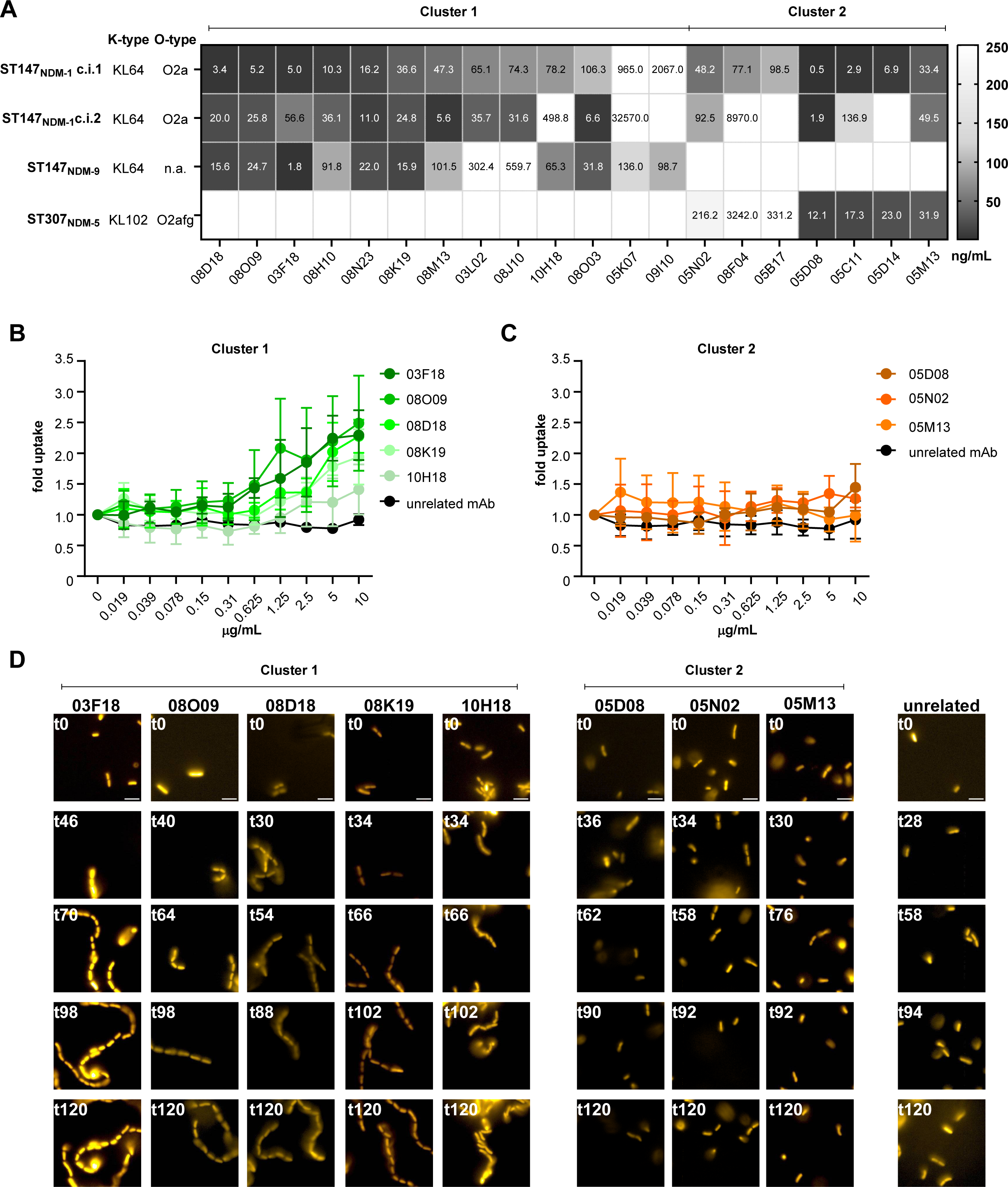
Functional *in vitro* characterization of bactericidal ST147_NDM-1_-targeting mAbs. (A) Heatmap reporting mean IC_50_ values resulting from F-SBA screening of 20 anti-Kp mAbs against the indicated Kp strains. Values for each mAb result from 2 to 6 replicated experiments. (B) Bacterial fold uptake by THP-1 macrophages measured in phagocytosis assay in the presence of anti-Kp mAbs from cluster 1 on ST147_NDM1_ c.i. 1-sfmCherry. Two-way ANOVA test to compare each anti-Kp mAb to the unrelated mAb resulted in the following *P* values at the minimum effective dose: p=0.0004 and p=0.001 for 03F18 and 08O09, respectively, at 0.625 µg/mL; p=0.016 for 08D18 at 1.25 µg/mL; p=0.004 and p=0.035 for 08K19 and 10H18, respectively, at 2.5 µg/mL. Experimental replicates: N=6 for 03F18; N=4 for 08O09; N=5 for 08D18; N=4 for 08K19; N=6 for 10H18. (C) Bacterial fold uptake by THP-1 cells measured in samples treated with anti-Kp mAbs from cluster 2 on ST147_NDM1_ c.i. 1-sfmCherry. Two-way ANOVA test to compare each anti-Kp mAb to the unrelated mAb resulted in no significant *P* values at any tested dose. Experimental replicates: N=2 for 05D08 and 05N02; N=3 for 05M13. (D) Panels show still frames from time-lapse imaging of on ST147_NDM1_ c.i. 1 bacteria expressing sfmCherry and incubated with 100 µg/mL of indicated mAbs for 120 min. Scale bar 5 µm.

Since phagocytes play a crucial role in Kp clearance during infection, mAbs displaying the highest bactericidal potency were then evaluated in macrophage-mediated opsonophagocytosis assays. Superfolder (sf) mCherry-overexpressing recombinant ST147_NDM-1_ was opsonized with escalating doses of mAbs prior the incubation with differentiated THP-1 cells. Then, fluorescence of internalized Kp was measured as a readout of mAb ability to promote opsonophagocytosis. Anti-KL64 mAbs were found to significantly increase Kp uptake by THP-1 cells at doses greater than 0.6 μg/mL, except for 10H18 which showed a very mild effect (**Figure 4B**). However, no significant Kp uptake was detected in the presence of anti-O-antigen mAbs at none of the tested doses (**Figure 4C**).

Finally, to analyze the impact of bactericidal mAbs on Kp cultures over time, time-lapse imaging of ST147_NDM-1_-sfmCherry incubated with increasing doses of mAbs was performed. Cluster 1 mAbs induced enchained bacterial growth in the 10 to 100 μg/mL range of concentrations (**Figure 4D**). Mechanistically, this phenotype was associated with the appearance of concatenated replicating bacteria which failed to generate separate sister cells (**Supplementary Video 1**). On the contrary, cluster 2 mAbs did not induce enchained growth at any tested dose (**Figure 4D; Supplementary Video 2**).

Overall, this multi-functional analysis indicated that isolated mAbs had an extremely potent bactericidal activity and that their functionality extended beyond opsonophagocytosis and implicated other effector mechanisms, such as enchained Kp growth. However, surprisingly, only anti-capsular mAbs were endowed with such poly-functionality.

### mAb poly-functionality is required for protection against fulminant ST147_NDM-1_ bloodstream infection *in vivo*

To address the fundamental question of whether picomolar bactericidal activity on its own correlates with mAb efficacy *in vivo*, or whether poly-functionality is required, the protective properties of the most potent mAb candidates (anti-capsular 08O09, and anti O-antigen 05D08 and 05N02) were evaluated in prophylaxis (PRO) and treatment (TRT) studies in an immunocompetent murine septicemia model^51^. A fulminant bloodstream infection model was selected to reflect the clinical history of donors from whom mAbs were isolated and was established to achieve ≥ 90% mortality within 24 hours of bacterial challenge in absence of therapy (**Figure 5A**). A relatively high bacterial inoculum (10^6.5^ CFU/mL) was selected to better mimic the disease severity in a clinical setting.

**Figure 5.**
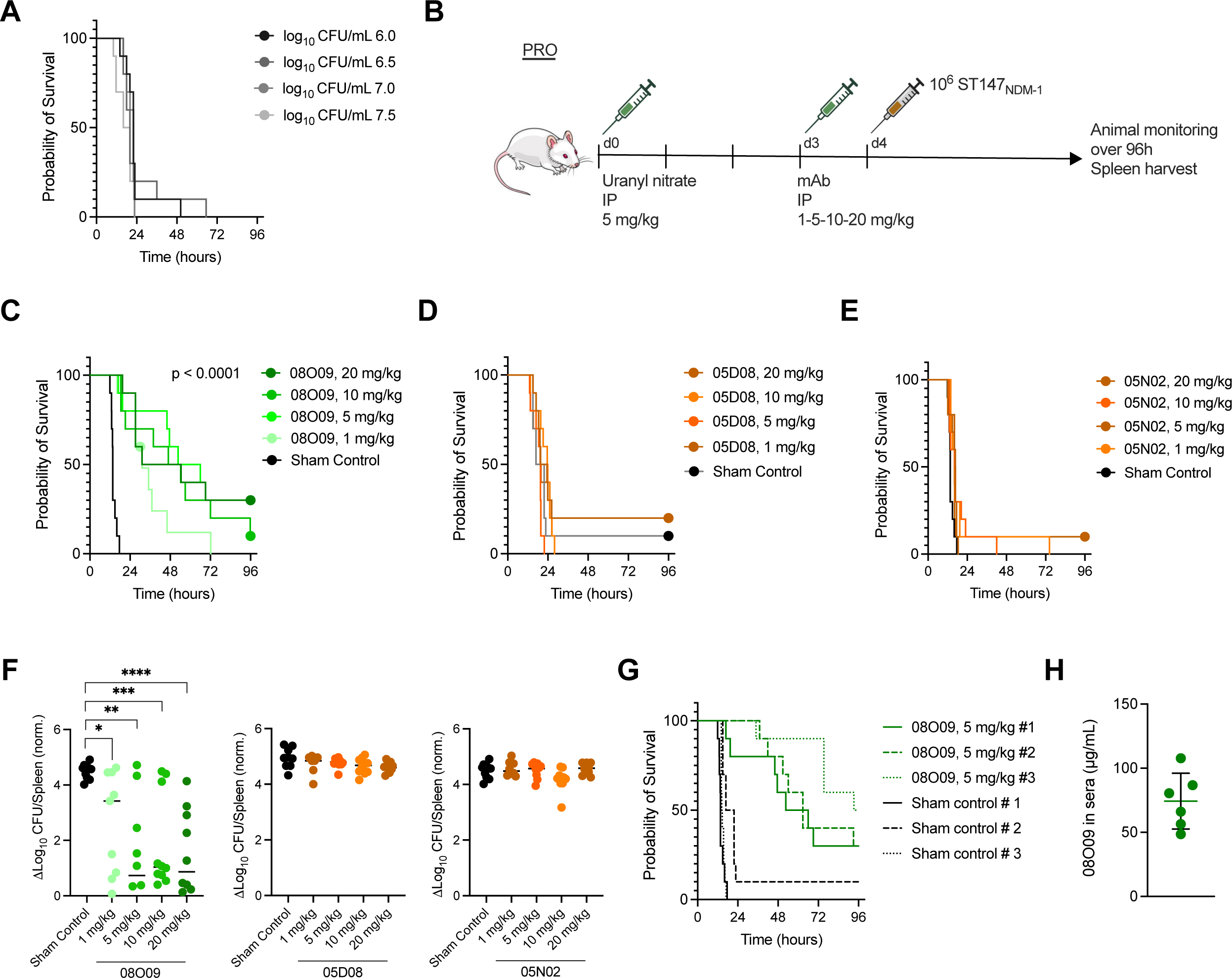
Evaluation of *in vivo* prophylactic efficacy of 08O09, 05D08 and 05N02 mAbs in immunocompetent ST147_NDM-1_ bacteremia model. (A) Survival analysis for groups of mice infected with ST147_NDM-1_ c.i. 1 at four tested inoculums (n=10 animals per group). (B) Scheme of the prophylaxis (PRO) study. (C-D) Survival analysis of 08O09 (C), 05N02 (D) and 05D08 (E) PRO regimens versus sham controls (n=10 mice per group). Indicated *P* value was calculated by log-rank test and refers to a comparison of every 08O09-treated group versus the control group for each indicated administration dose. (F) Spleen load of ST147_NDM1_ in 08O09 PRO study. Log_10_ CFU/spleen at endpoint (n=10 animals per group) was normalized to non-infected controls (n=6 animals). Indicated are mean values; Mann-Whitney test * p< 0.05, ** p< 0.01, *** p<0.001, **** p<0.0001. (G) Survival analysis of 08O09 PRO regimens administered as a single dose of 5 mg/kg in three independent experiments versus sham controls (n=10 mice per group). Log-rank test to compare each 08O09-treated group versus the respective control group resulted in the following *P* values: <0.0001 for study #1; 0.0049 for study #2, <0.0001 for study #3. (H) Concentration of 08O09 mAb in mouse plasma was measured by ELISA 24 hours after 5 mg/kg i.p. administration (n=6 animals). Statistical bar indicates mean values ± SD.

To assess the *in vivo* PRO activity against ST147_NDM-1_, mice were administered with 1, 5, 10, and 20 mg/kg mAb dose 24 hours prior to bacterial inoculation (**Figure 5B**). 08O09 PRO administration at all four examined doses prolonged survival compared with sham control, with 5 mg/kg and higher doses showing comparable survivals (**Figure 5C**). Instead, 05D08 and 05N02 mAbs did not prolong animal survival, at none of the administered doses (**Figure 5D and E**). Consistent with this observation, CFU counts in the spleen showed a reduction upon treatment with 08O09 (mean reduction of 1.9 to 3 log_10_ CFU) but did not show reduction in case of 05D08 and 05N02 (**Figure 5F**). PRO experiment with 5 mg/kg 08O09 dose was repeated on 30 animals in total in three separate experiments, showing consistency and reproducibility, with approximately 50% survival up to 96 hours (**Figure 5G**). Plasma concentration of 08O09 associated with *in vivo* efficacy amounted to 74.3 ± 21.7 µg/mL (**Figure 5H**). Hence, consistent survival results and decrease in organ bacterial load using 08O09 5 mg/kg PRO across runs demonstrated the robustness of the model to delineate efficacy *in vivo*.

Considering these results, the treatment (THR) efficacy of 08O09 administered intravenously at 1, 5, 10, and 20 mg/kg one-hour post-bacterial challenge was assessed **(Figure 6A)**. In this case, 08O09 THR at the four examined doses resulted in significantly prolonged survival compared with sham control **(Figure 6B)**, which was associated with 1 to 2.4 mean log_10_ CFU reduction in the spleen **(Figure 6C)**. Survival was comparable among groups treated with 5 mg/kg and higher doses.

**Figure 6.**
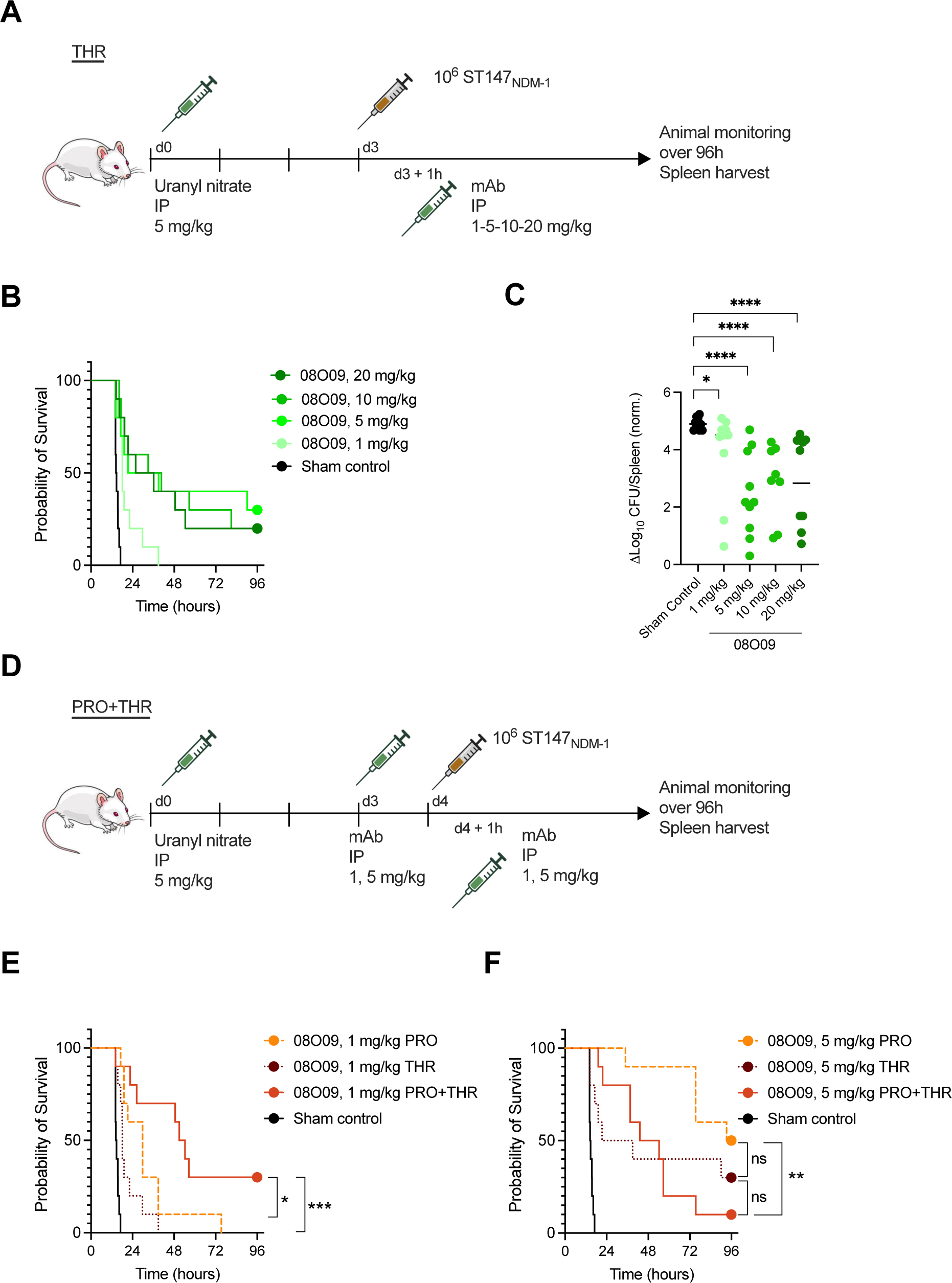
Evaluation of *in vivo* therapeutic and combined efficacy of 08O09, 05D08 and 05N02 mAbs in immunocompetent ST147_NDM-1_ bacteremia model. (A) Scheme of treatment (THR) study. (B) Survival analysis of 08O09 THR regimens versus sham controls (n=10 mice per group). Log-rank test to compare each 08O09-treated group versus the respective control group resulted in the following *P* values: 0.0002 for 1 mg/kg; 0.0005 for 5 mg/kg, and <0.0001 for 10 and 20 mg/kg. (C) Spleen load of ST147_NDM1_ in 08O09 THR study. Log_10_ CFU/spleen at endpoint (n=10 animals per group) was normalized to non-infected controls (n=6 animals). Mann-Whitney test * p< 0.05, **** p<0.0001. (D) Scheme of prophylaxis plus treatment (PRO+THR) study. (E) Survival analysis of 08O09 PRO regimens and PRO+THR regimens at 1 mg/kg doses (n=10 mice per group). Indicated *P* values were calculated by log-rank test and refer to a comparison of PRO+THR group versus either PRO (p<0.05) or THR group (p<0.001). (F) Survival analysis of 08O09 PRO regimens and PRO+THR regimens at 5 mg/kg doses (n=10 mice per group). Indicated *P* values were calculated by log-rank test and refer to comparisons indicated by brackets.

Lastly, the protective properties of 08O09 in a combined prophylaxis/treatment (PRO+THR) regimen were measured. In this case, the first dose of 08O09 was administered intraperitoneally 24 hours before bacterial inoculation, and the second dose was administered intravenously one-hour post-bacterial challenge **(Figure 6D)**. Interestingly, an advantage of using a combinatorial treatment, with 30% survival till 96 h, was observed only for 1 mg/kg dosage that was otherwise suboptimal in monotherapy regimens (**Figure 6E**). However, when 08O09 was administered at 5 mg/kg, PRO monotherapy performed significantly better than combinatorial treatment (**Figure 6F**), suggesting that while a suboptimal mAb dosing could be compensated by combined regimen, an optimal mAb dosing may perform well as a monotherapy. Overall, these experiments demonstrated that an anti-KL64-specific poly-functional mAb protects animals in PRO and THR regimens by efficiently reducing the systemic load of pandrug-resistant ST147_NDM-1_.

## DISCUSSION

In this work, we presented a rationally designed pipeline to screen and isolate protective human mAbs against hypervirulent and pandrug-resistant ST147_NDM-1_ Kp. This strain, which was obtained from a highly persistent nosocomial outbreak that started in 2018 in the hospital of Pisa in Italy, is now endemic in the region^19–21^. Moreover, ST147_NDM-1_ Kp is also spreading globally with high speed and has already been described in Europe^20,52^, USA^53^, and Southeast Asia^54^. Our antigen-agnostic approach resulted in the selection of human mAbs that can be exploited as therapeutic and prophylactic agents against ST147_NDM-1_. Of note, we identified *in vitro* assays that correlate with protection. Through this strategy, we discovered and characterized the first human mAb which interacts with capsule type KL64 and protects against fulminant bloodstream infection by the pandrug-resistant Kp strain.

Our data highlight the importance of developing robust predictive assays to correlate *in vitro* mAb functionality with *in vivo* efficacy. Out of more than 200 ST147_NDM1_-specific mAbs isolated from convalescent donors, we found 20 highly bactericidal antibodies targeting either the capsule or the O-antigen, all working in the picomolar range of concentrations. However, despite the similar potency in the bactericidal assay, only a subset of them promoted phagocytosis and bacterial enchained growth, which correlated with protection against bloodstream infection. This evidence indicated that bactericidal activity, which is usually a strong correlate of protection, is not sufficient for *in vitro* mAbs characterization and identification of the best candidate, whereas *in vitro* poly-functionality is an essential feature that protective mAbs must bear, particularly in the case of those targeting the pandrug-resistant strain ST147_NDM-1_. Lack of correlation with complement-mediated bactericidal activity and the positive correlation with phagocytosis are aligned with the knowledge that Kp killing and clearance is impaired when cellular innate immune functionality is decreased or deficient^55,56^. In particular, increased susceptibility to Kp infections has been observed in association with neutrophil deficiencies^56,57^.

Noteworthy, the capsule is an important virulence factor for Kp pathogenesis which shields bacteria against the host immune response^58–61^. Previous studies showed that mutations preventing capsule synthesis transformed Kp strains into avirulent bacteria when infecting rats or mice^59,62,63^. Likewise, it has been reported that in some cases anti-O-antigen mAbs were protective only against non-encapsulated Kp strains^64^, suggesting that Kp capsule is an important immune evasion factor. Although we did not explore this aspect in the present work, we found that O-antigen-specific mAbs were devoid of *in vitro* poly-functionality and *in vivo* efficacy, implying that KL64 capsule might limit their access to the cognate epitopes thereby interfering with their action. Another possibility that has remained unexplored is that, during *in vivo* infection, KL64 capsule production could be dynamically modulated to further increase the shielding effect. Hence, KL64 capsule represents a strategically important immunodominant antigen to be neutralized.

By correlating *in vitro* and *in vivo* results we propose that anti-KL64 capsule mAbs protect *in vivo* by means of two main mechanisms of action, the first being macrophage-mediated killing. An additional underexplored mechanism of immune protection employed by KL64-targeting mAbs is triggering bacterial enchained growth *in vitro*. Bacterial concatenation induced by secretory dimeric IgA (sIgA) has been previously described as a protective mechanism restricting *Salmonella* Typhimurium growth and facilitating its clearance from the intestine^65,66^. Interestingly, here we observed that this phenomenon applies to Kp as well and can be promoted by monomeric IgG. Moreover, considering that only anti-capsule and not anti-O-antigen mAbs induce enchained growth, we ought to think that targeting the most external bacterial layer (*i.e.* the capsule) is essential to promote enchainment.

ST147_NDM-1_ has been classified as a pandrug-resistant and hypervirulent strain^26^ which has not only been responsible for local outbreaks in Italy^19–21^ but has been causative of infections worldwide^20,52–54^. Additionally, genomic surveillance reports indicate that KL64, associated with enhanced virulence^67^, is replacing some previously prevalent capsular loci through intraspecies clone shifting^68^ and became the third emerging capsule type for hypervirulent strains after K1 and K2 serotypes^4,69^. Indeed, carbapenem-resistant KL64-expressing ST11 Kp represents the most common AMR lineage in China^70^. Taken together, these data underline the importance of anti-capsular mAbs to contain and prevent future outbreaks caused by ST147_NDM-1_ and other KL64-bearing Kp isolates. As such, 08O09 mAb, developed in the pursuit of an effective therapy against a regional outbreak of a pandrug-resistant strain, might represent an excellent candidate for treatment and prophylaxis of KL64 Kp infections globally.

## LIMITATIONS OF THE STUDY

Hyper-capsulation of ST147_NDM-1_ Kp may explain why, despite applying an antigen-agnostic approach, we pinpointed only anti-capsule and anti-OAg bactericidal mAbs, while no bactericidal antibodies against other type of antigens were identified. Indeed, Kp protein antigens may be masked by the thick OAg and capsule layers, which probably elicited most of the humoral immune response with bactericidal activity. While the work resulted in the identification of a candidate effective medication against ST147_NDM-1_ Kp, clinical application of 08O09 to Kp infections will be limited to the KL64 serotype. In the framework of exploiting mAbs for anti-Kp universal vaccine design, future research efforts should be directed towards the identification of functional antibodies against conserved targets. To this end, implementation of specific screening strategies will be instrumental. Finally, careful choice of the target patients who may benefit from mAb-based therapy should be made since ST147_NDM-1_ Kp can cause acute infections as well as chronic colonization which may require differential treatment protocols.

## Supporting information

SupplementalVideo1_Cluster1 mAb

SupplementalVideo2_Cluster2 mAb

## ACKNOWLEDGMENTS

We wish to thank all patients who participated in this study. We acknowledge Federica Romano, Giulia Fantoni, Giulio Pierleoni and the Clinical Studies team at Fondazione Toscana Life Sciences for technical support and help with clinical study documentation; Valentina Galfo (University of Pisa) for the contribution to patient sampling and enrolment; Simona Tavarini and Chiara Sammicheli (GlaxoSmithKline, Siena, Italy) for help with cell sorting and flow cytometry; Maria Grazia Pizza (Imperial College, London, UK) for insightful discussion; Francesca Micoli, Carlo Giannelli, Omar Rossi, Gianmarco Gasperini (GlaxoSmithKline, Siena, Italy) for technical advice on Klebsiella antigen purification; Chris Whitfield and Steven Kelly (Department of Molecular and Cellular Biology, University of Guelph, Guelph, Ontario, Canada) for the generous gift of Kp LPS; Maria Grazia Cusi (University Hospital of Siena), Deborah Hung (The Broad Institute of MIT and Harvard, Cambridge, MA, USA), Joshua Liang Chao L.C. Wong and Gad Frankel (Department of Life Sciences, MRC Centre for Molecular Bacteriology and Infection, Imperial College London, London, UK) for providing Kp strains and insightful discussion.

## AUTHOR CONTRIBUTIONS

Conceptualization: CS, AK, RR; methodology: ER, VZGF, SSB, IP, GM, AK; investigation: ER, VZGF, SSB, IP, GM, GB, SS, GC, SY, LC, MR, MT, NM, CM, CDS, AC, LC,; bioinformatic analysis: GM, DC; resources: GT, CG, LC, SB, FM, MF, DPN, KA; data curation: GM, DC, MF, DL; writing – original draft: ER, VZGF, SSB, GM, GB; writing – review & editing: ER, VZGF, SSB, GM, DPN, KA, CS, AK, RR; visualization: ER, VZGF, SSB, GM, AK; supervision and funding acquisition: CS, AK, RR.

## FUNDING

This project was supported by funds of the Centro Regionale per la Medicina di Precisione (CREMEP) and by PROREACT (progetto finanziato con le risorse del Fondo per il rilancio degli investimenti delle amministrazioni centrali dello Stato e allo sviluppo del Paese, di cui all’art. 1 comma 95, n. 145/2018, fondi 2019-2023, del Ministero della Salute).

## DECLARATION OF INTERESTS

Patent application 102023000000924 describing isolated mAbs has been filed by Fondazione Toscana Life Sciences.

R.R. hold shares in the GSK group of companies and declare no other financial and non-financial relationships and activities. Remaining authors declare no competing interests.

## MATERIALS AND METHODS

### Enrollment of Kp-infected patients

Clinical protocol Klebsiella01 was approved by the ethics committee of the University Hospital of Pisa (Parere 16526). All enrolled patients signed a written informed consent form before taking part in the study which was conducted in accordance with Good Clinical Practice (GCP) guidelines and ethical principles of the Declaration of Helsinki. Patients’ information can be found in Table S1.

### Bacterial strains and culture conditions

Bacterial strains used in this work are listed in Table 1. Kp isolates were grown at 37°C in LB broth in the presence of the appropriate antibiotic (5 μg/mL meropenem for ST147_NDM-1_ c.i.1, ST147_NDM-1_ c.i.2, ST147_NDM-9_,ST307_NDM-5_, ST13_OXA-48_, ST512_VIM_; 150 μg/mL hygromycin for sfmCherry ST147_NDM-1_ c.i. 1), or on LB-agar plates supplemented with the appropriate antibiotic (5 μg/mL meropenem for ST147_NDM-1_ c.i.1, ST147_NDM-1_ c.i.2, ST147_NDM-9_,ST307_NDM-5_, ST13_OXA-48_, ST512_VIM_; 150 μg/mL hygromycin for sfmCherry ST147_NDM-1_ c.i. 1). All other bacterial strains were grown in LB.

### Single-cell sorting of memory B cells from convalescent donors

Peripheral blood mononuclear cells (PBMCs) were isolated from heparin-treated whole blood by density gradient centrifugation (Ficoll-Paque PREMIUM, GE Healthcare). After separation, PBMC were stained with Live/Dead FSV780(BD Horizon) in 100 µL final volume diluted 1:1000 at room temperature (RT). After 20 min of incubation, cells were washed with phosphate-buffered saline 1X (PBS) and unspecific bindings were saturated with 50 µL of 20% normal rabbit serum (Life Technologies) in PBS. Following incubation at 4°C for 30 min, cells were washed with PBS 1X, centrifuged at 1200 rpm for 8 min, and stained with CD19 APC (BD cat# 561742), IgM PeCF594 (BD cat# 562539), CD27 APCR700 (BD cat# 565116), IgD PE (BD cat# 562024), CD3 PE-Cy7 (BD cat# 560910), CD14 PE-Cy7 (BD cat# 560919) and CD56 PECy7 (BD cat# 560916) in Staining Buffer (1% Fetal Bovine Serum in PBS) at 4°C for 30 min. Following additional washing, cells were resuspended in Sorting Buffer (PBS/EDTA 2.5 mM). Stained memory B cells (MBCs; CD3/CD14/CD56^-^ CD19^+^ CD27^+^ IgD^-^ IgM^-^) were single-cell sorted with a BD FACSAria™ Fusion (BD Biosciences) into 384-well microplates containing 3T3-CD40L feeder cells and were incubated with IL-2 and IL-21 for 14 days^71^.

### ELISA screening of mAb binding against *Klebsiella pneumoniae*

Supernatants resulting from single-cell sorting of MBCs were used as a source of IgG1 and IgA monoclonal antibodies (mAbs) to test their binding to whole bacteria in a high-throughput enzyme-linked immunosorbent assay (ELISA). Pools of ST147_NDM-1_ c.i. 1 and ST147_NDM-1_ c.i. 2 (**Table 1**) were grown in LB medium, pelleted at exponential phase (OD_600_=0.5), resuspended in PBS, and plated into 384-well plastic plates (Greiner, ref. 781101). Plated bacteria were fixed at RT for 30 min in 0.5 % of paraformaldehyde. Plates were then blocked in PBS plus 2% Calf Serum (Euroclone, ECS0040L) at RT for 1 h. After incubation, bacteria were incubated for 1 h with MBC supernatants diluted in PBS plus 2% Calf Serum. Next, anti-human IgG and anti-human IgA secondary antibodies conjugated with Alkaline Phosphatase (Southern Biotech) were added for 45 min. To detect bacteria-mAbs binding, pNPP (p-nitrophenyl phosphate; Sigma-Aldrich) was used as soluble substrate and the final reaction was quantified at a wavelength of 405 nm by using the Varioskan Lux Reader (Thermo Fisher Scientific). After each incubation step, plates were washed three times with 100 µl per well of washing buffer (PBS and 0.05% Tween-20). Sample buffer was used as a blank and we considered as positive hits those wells showing an OD_405_ value of at least 2-fold superior to blank. Cells corresponding to positive hits were lysed in 25 μL of a buffer containing RNAse Out 0.2 U/µL, ultrapure BSA 1 mg/mL and DEPC H_2_O (Thermo Fisher Scientific) and stored at −80°C until use.

### Single-cell RT-PCR and nested PCR were used to amplify V_H_ and V_L_

cDNA was synthesized from 5 μL of MBC lysates. Reverse transcription (RT) reaction was performed by adding 25 μL per well of a mix containing 1 μL of random hexamer primers (50 ng/mL),1 μL of dNTPs (10 mM), 2 μL 0.1 M DTT, 40 U/μL Rnase OUT, MgCl_2_ (25 mM), 5 μL of 5X buffer, 0.25 µL of Superscript IV reverse transcriptase (Invitrogen) and nuclease-free water (DEPC) and RT-PCR conditions were 42°C/10 min, 25°C/10 min, 50°C/ 60 min and 94°C/5 min.

After cDNA synthesis, two additional rounds of PCR were performed to obtain the variable regions of the heavy (V_H_) and light (V_L_) chains. Briefly, in the first round of PCR (PCR I) a total volume of 25 μL containing 4 μL of cDNA, 10 μM of VH or 10 μM VL/VK primer mix (**Table S2**), 0.5 μL of dNTPs (10 mM), 1.5 μL MgCl_2_ (25 mM), 5 μL of 5X Kapa Long Range Buffer, and 0.125 μL of Kapa Long Range Polymerase (Sigma) was added in each well and amplified using the following conditions: 95°C/3’, 5 cycles at 95°C/30’’, 57°C/30’’, 72°C/ 30’’ and 30 cycles at 95°C/30’’, 60°C/30’’, 72°C/30’’ and 72°C/2’. 3 μL of un-purified PCR I products were used as a template for the nested PCR (PCR II) using the same cycling conditions and primers indicated in **Table S2**. PCR II products were then purified by Millipore MultiScreen PCR 96 plate according to the manufacturer’s instructions. Samples were eluted in 30 μL nuclease-free water pre-warmed at 50°C and quantified by NanoDrop One (Thermo Fisher Scientific).

### Cloning of V_H_ and V_L_ and recombinant antibody expression by TAP transfection

Amplified antibody sequences were ligated into human IgG1 expression vectors and used to produce transcriptionally active PCR (TAP) products. Briefly, IgG1, Igκ or Igλ expression vectors were digested with AgeI, SalI and XhoI restriction enzymes, respectively. 25 ng of linearised plasmids were ligated with 75 ng of purified V_H_ and V_L_ by Gibson Assembly (New England BioLabs). The reaction was performed in a final volume of 5 μL. The ligation product was 10-fold diluted in DEPC water and used as a template for transcriptionally active PCR (TAP) reaction which allowed the direct use of linear DNA fragments for *in vitro* expression. Functional promoter (human CMV) and terminator sequences (SV40) were attached onto PCR II products directly by amplification using specific primers (**Table S2**). TAP-PCR was performed in a total volume of 25 μL containing 0.25 μL of Q5 polymerase (NEB), 5 μL of GC Enhancer (NEB), 5 μL of 5X buffer, 0.5 μL dNTPs (10 mM), 0.125 μL of forward/reverse primers and 3 μL of ligation product, and using the following conditions: one step of 98°C/2’, 35 cycles of 98°C/10’’, 61°C/20’’, 72°C/1’ and an extension of 72°C/5’. Once purified and quantified, TAP products were transfected into Expi293F cells to allow small scale production of recombinant mAbs. Briefly, cells were transfected with both TAP amplifications in a final volume of 1 ml into 96 deep well plate (Eppendorf), and after 7 days of expression mAb-containing supernatants were harvested by centrifugation.

### Quantification of TAP-produced mAbs by ELISA

To quantify the concentration of mAbs in each supernatant, ELISA plates were coated with 2 µg/ml of goat anti-human IgG (Southern Biotech) at 4°C overnight. Plates were washed 3 times with PBS plus Tween-20 0.05 %, blocked in PBS plus BSA 1% at 37°C for 1 h. Then samples were washed and incubated with TAP-produced mAbs diluted in sample buffer (PBS – BSA 1 %-Tween-20 0.05%) at 37°C for 1 h. Following additional washings and incubation at 37°C for 1 h with AP-conjugated anti-goat IgG secondary antibody (Southern Biotech), absorbance was read by addition of PNPP. Concentrations were evaluated by linear regression analysis built by potting OD_405_ values against a standard curve generated by titration of a human IgG-unlabeled antibody (Southern Biotech).

### Large scale expression and purification of mAbs

Expi293F cells (Thermo Fisher) were transiently transfected with plasmids carrying the heavy and the light chains of each antibody with a 1:2 weight/weight ratio. Cells were grown for six days at 37°C with 8% CO_2_ and 125 rpm shaking, with an optimized cocktail of enhancers 1 and 2 (Thermo Fisher) added on day 1 post-transfection. Two mAb harvests were performed on the third and sixth day by pelleting the cells at 1,100 x g for 10 min at RT and supernatants were pooled and clarified by centrifugation (3,000 x g for 15 min at 4°C), followed by 0.45 mm filtration. mAbs were purified at RT by affinity chromatography on the AKTAgo purification system (Cytiva) using the HiTrap Protein G HP column (Cytiva), which binds to the Fc domain. Specifically, the column was equilibrated in 0.02 M sodium phosphate buffer pH 7 at a flow rate of 1 mL/min, which was used also for the following steps. After sample injection, the column was washed with 10 column volumes (CV), followed by mAb elution with 10 CV of 0.1 M glycine-HCl, pH 2.7. mAb pool was dialyzed in PBS pH 7.4 using Slide-A-Lyzer G2 Dialysis Cassette 3.5K (Thermo Scientific) overnight at 4°C. For each purified antibody, the concentration was determined by measuring absorbance at 280 nm by Nanodrop. All purified antibodies were aliquoted and stored at −80°C.

### Serum Bactericidal Assay (SBA)

Two days prior to the assay, glycerol stocks of Kp were streaked on LB-Agar plates and incubated overnight (ON) at 37°C. On the following day, a single colony was inoculated in 4 mL of LB and incubated ON at 37°C. Cultures were expanded to 10 mL LB in a 125 mL flask to obtain an OD_600_ of 0.05 and incubated at 37°C with shaking until the exponential phase (OD_600_ 0.4-0.6) was reached. Bacteria were then 1:10 diluted in PBS.

Appropriate baby rabbit complement (BRC, Cedarlane) concentrations to be used in the assay were established through complement sensitivity tests. Briefly, 2×10^5^ bacteria resuspended in PBS were seeded in round bottom 96-well plates, and BRC concentrations were screened starting from 50% in 11 serial 2-fold step dilutions. After 2 h incubation at 37°C, bacteria were pelleted by centrifugation. The supernatant was discarded, and bacteria were resuspended in 30 µl of PBS and transferred into a White Optiplate (Perkin Elmer). 30 µl of BacTiter-Glo™ 1X (Promega) was added to each well and luminescence was measured by the Varioskan Lux microplate reader (Thermo Fisher Scientific) with an exposure time of 500 ms. Data were plotted and analysed using GraphPad Prism. 12.5 % BRC was chosen as the BRC concentration allowing to achieve complement-based antibody killing without significant toxicity.

For luminescence-based SBA (L-SBA) with TAP-expressed mAbs, 2×10^6^ bacteria/mL of ST147_NDM-1_ were seeded into a 96-well U-round bottom plate in the presence of 12.5% BRC, with the addition of four serial dilutions of TAP-expressed mAbs (1:10, 1:50: 1:250: 1:1250), in a total volume of 100 µl per well. Upon 2 h incubation at 37°C, bacteria were pelleted by centrifugation at 4000 xg. Bacterial pellets were resuspended in 30 µl of PBS per well and transferred into a white Optiplate (Perkin Elmer). 30 µl of BacTiter-Glo™ 1X (Promega) was added to each well and luminescence was measured by the Varioskan Lux microplate reader (Thermo Fisher Scientific) with 500 ms exposure. In each experiment, luminescence values obtained were used to calculate the median value for each dilution factor. The difference between the luminescence signal of each mAb and the median value was measured and plotted as percentage with 30 % cut-off. For fluorescence-based SBA (F-SBA), 2×10^6^ bacteria/mL in PBS were seeded in 384-well black clear-bottom plate (ViewPlate®-384 F TC, PerkinElmer) in the presence of 12.5 % BRC, in a final volume of 50 µl per well. Purified recombinant mAbs were added in a 3-fold step serial dilution panel. After 2 h incubation at 37°C, 40 µl LB and 10 µl of 0.025% resazurin (Sigma-Aldrich) were added to each well. Fluorescence (λ_Ex_ 560 nm and λ_Em_ 590 nm, exposure: 250 ms) was measured by the Varioskan Lux microplate reader (Thermo Fisher Scientific) upon 2 h incubation at 37°C. GraphPad Prism was used to plot and analyse data, as well as to extrapolate IC_50_ values.

### Flow cytometry analysis of mAb binding to bacterial surface

Binding of mAbs to the bacterial strains listed in Table 1 was assessed in the exponential phase of growth in LB (OD_600_ 0.4-0.6). Bacteria were pelleted and resuspended in an equal volume of PBS-BSA 1%. 100 μL of bacteria were plated in each well of a round bottom 96-well plates by centrifuging at 4000 x g for 5 min, followed by incubation with 5 µg/ml of purified antibodies at RT for 1 h. Plates were centrifuged, and pellets were washed three times with PBS – BSA 1% and incubated at RT for 30 min in the dark with 1:2000 diluted Alexa488-conjugated α-human IgG secondary antibody. After an additional centrifugation and washing step, bacteria were fixed with 0.5% of paraformaldehyde at RT for 30 min, washed again, resuspended in PBS – BSA 1% to an OD_600_ of 0.05 and read by their fluorescence. Samples were acquired on the BD FACS Canto II (BD Biosciences, USA). Data were analysed by FlowJo software v10 (BD Biosciences, USA).

### Antibody sequence analysis

Immunoglobulin genes were identified with NCBI IgBlast^72^ and named according to the IMGT nomenclature (in the time between 2020 and 2022). IGHV gene somatic hypermutations were counted from the start of FWR1 until the end of FWR3. Insertions or deletions were counted as one single mutation.

### Genomic analysis of Kp isolates

ST147 strains were sequenced using both short- and long-reads technology. High-throughput-sequencing was performed on the MiSeq platform (Illumina; San Diego, CA, USA) with a paired-end layout of 150 bp. Paired-end short reads were quality-checked and poor-quality reads were filtered using fastp v0.20.1^73^. The long-read library was prepared with multiplexing and sequenced according to the manufactures’ guide using flow cell R9.4.1 (Nanopore). The quality of long reads was controlled by being mapped with their corresponding short reads using Filtlong v0.2.0 (https://github.com/rrwick/Filtlong) with minimum quality and length as Q8 and 2,000 bp respectively. Along with the corresponding clean reads, long reads were fed into the hybrid assembler Unicycler v0.4.8^74^ and run under the conservative mode. ST and capsule typing prediction was performed using Kleborate v2.0.0^10^ and Kaptive^75^. Roary^76^ version 3.13.0 was used for pan-genome analysis, and the core gene alignment from Roary was used to generate a core gene single nucleotide polymorphism (SNP) tree. First, SNP sites in the core gene alignment from Roary were concatenated with snp-sites^77^ version 2.4.1. The resulting concatenated SNPs were used in IQ-Tree^78^ version 1.6.8 to create a maximum likelihood (ML) tree. The optimal evolutionary model was selected by using ModelFinder plus. Raw sequencing data have been released under BioProject PRJNA1067287.

### Immunoblot analysis of mAb binding to Kp lysates

For sample preparation, bacteria from glycerol stocks were grown overnight on LB-agar plates. Single colonies of Kp were picked from plates and grown overnight with static incubation at 37°C in LB. On the following day, bacteria were grown at 37°C starting from OD_600_ 0.05 with shaking in LB until they reached the exponential phase (OD_600_ 0.4-0.6). Polysaccharide extracts were prepared by phenol extraction. Following overnight growth, bacteria were pelleted and resuspended in PBS. Upon 1 h incubation at 100°C, an equal volume of phenol was added, and samples were incubated at 70°C for 1.5 h. Samples were centrifuged at 20000 xg at 4°C for 30 min, then the water phase was recovered. SDS-PAGE samples were prepared by adding 1:2 tris-glycine loading buffer and incubated for 2 min at 85°C. SDS-PAGE gel was transferred onto PVDF membrane using the iBlot™ Gel Transfer Device and Stacks (Thermo Fisher). Membranes were blocked for 1h at room temperature in TBS 1x/0.1% Tween-20/5% milk. Purified mAbs were used at 1µg/mL in TBS 1x/0.1% Tween-20/5% milk. Following an overnight incubation at 4°C, membranes were washed 3 times with TBS 1x/0.1% Tween-20. Incubation with secondary antibody (goat anti-human Fab) diluted 1:75000 in TBS 1x/0.1% Tween-20/5% milk was carried out for 1 h at room temperature. Membranes were washed 3 times in TBS 1x/0.1% Tween-20 and then developed with chemiluminescence readout.

### Characterization of mAb binding by high-resolution and high-content confocal microscopy

A single colony of bacteria was picked from an LB-agar plate and grown ON in LB with 5 μg/mL meropenem. ST147_NDM1_ stably expressing super-folder(sf)mCherry were grown in LB with 150 μg/mL hygromycin. The overnight culture was diluted in LB without antibiotics to OD_600_ 0.025 or 0.05 and plated into a 96-well Phenoplate (Perkin Elmer, 6055300). Upon incubation for 2 h at 37°C without CO_2_ in static conditions, adherent bacteria were fixed in 4% paraformaldehyde (PFA)/PBS (Thermo Fisher) or Cytofix (BD) for 15 min at RT. Alternatively, overnight grown bacteria were re-launched in LB with antibiotics until exponential phase (OD_600_ 0.4-0.6). Then, bacteria were pelleted, resuspended in PBS at OD_600_ 0.25 and transferred into a 96-well Phenoplate. Finally, samples were briefly spinned at the bottom of the plate before fixation in 4% PFA.

For experiments aimed at staining a single anti-Kp mAb, selected antibodies were diluted in a solution of PBS/1% BSA at a concentration of 0.5 μg/mL and incubated for 30 min at RT. Then, samples were washed in PBS and a mixture of goat anti-Human Alexa488 conjugated secondary antibody (Thermo Fisher, diluted 1:2000) and DAPI (Thermo Fisher, diluted 1:2000) in PBS/BSA1% was added for 30 min at RT. Finally, samples were washed in PBS and 50 μL of 1% Low Melting (LM) agarose (Sigma) were distributed in each well. Samples were stored at 4°C and imaging was performed within the following 24 hours.

For experiments in which multiple anti-Kp mAbs were simultaneously stained, antibodies were previously conjugated with Alexa488, Alexa555 and Alexa647 fluorophores, using the Zip Alexa Fluor™ Rapid Antibody Labelling Kits (Thermo Fisher) following manufacturer instructions. After fixation, samples were first blocked in PBS/1% BSA for 30 min at RT and then incubated with a mixture of fluorophore-conjugated mAbs (each one at 1 μg/mL) and DAPI (1:2000) in PBS/1% BSA. Following 30 min incubation at RT, samples were washed in PBS and prepared for imaging by adding 50 μL of 1% LM agarose. Samples were imaged on the following day.

### Construction of sfmCherry-expressing recombinant Kp strain

The ST147_NDM-1_ strain expressing sfmCherry under control of the pTAC promoter was obtained as follows. Codon-optimized tandem copies of the sfmCherry gene fused to the pTAC promoter were synthesized by GeneArt (Thermo Fisher Scientific) and cloned into the HindIII and SpeI restriction sites of the pSEVA23a1 vector^79^ where a hygromycin resistance cassette had been inserted in BamHI and XbaI. Two copies of sfmCherry were employed to maximise expression of the fluorescent protein, according to what had been reported by Kolbe and colleagues^80^. Restriction enzymes were from New England Biolabs. ST147_NDM-1_ was transformed by electroporation with the resulting plasmid using an Eporator instrument (Eppendorf) and transformants were selected on LB agar plates containing 150 μg/mL hygromycin. Transformants were validated by colony PCR and expression of sfmCherry was verified by flow cytometry and confocal microscopy. One of the transformed colonies was saved for further analyses.

### Complementation of ST147_NDM-9_ strain

The O-antigen-negative phenotype of the ST147_NDM-9_ Kp strain was complemented as follows. The *wbbO* gene of ST147_NDM-1_ Kp, retrieved by whole genome sequencing, was synthesized by GeneArt (Thermo Fisher Scientific) and cloned, under control of the pTAC promoter, into the HindIII and SpeI restriction sites of the pSEVA23a1 vector^79^ where a hygromycin resistance cassette had been inserted in BamHI and XbaI (all restriction enzymes were from New England Biolabs). Correct cloning was confirmed by Sanger sequencing (Eurofins). ST147_NDM-9_ was transformed by electroporation with the resulting construct according to standard procedures using an Eporator instrument (Eppendorf). Transformants were selected on LB agar plates containing 150 μg/mL hygromycin. One of the selected colonies was named ST147_NDM-9_/pSEVA23a1_*wbbO* and saved for further analyses.

### Opsonophagocytosis assay

THP-1 cells (ATCC) were maintained in RPMI 1640 containing GlutaMax (Thermo Fisher) and complemented with 10 mM Hepes (Thermo Fisher), 1 mM Sodium Pyruvate (Thermo Fisher) and 10% fetal bovine serum (FBS) (Thermo Fisher). Three days before the assay, 50,000 cells per well were seeded into a 96-well black-shielded Optiplate (Perkin Elmer) in the presence of 20 ng/mL phorbol-12-myristate-13-acetate (PMA) to promote monocyte differentiation into macrophages. The day after, PMA was washed out and cells were maintained in fresh medium for two additional days.

Infection of macrophage was performed using ST147_NDM1_-sfmCherry expressing Kp. Bacteria were grown ON and re-launched the following morning to reach OD_600_ 0.5. After centrifugation, bacteria were resuspended in phagocytosis media (RPMI 1640+GlutaMax, 10 mM Hepes, 1 mM Sodium Pyruvate) to have a 1:2 dilution of the initial culture volume. To allow bacterial opsonization, mAbs dilutions were prepared in 25 μL of phagocytosis media and incubated with 25 μL of bacteria for 30 min at 37°C in shaking conditions (500 rpm). This mixture was then added to differentiated macrophages and centrifuged for 3 min at room temperature to synchronize the infection. Following the incubation of 1 hour at 37°C in the presence of 5% CO_2_, samples were treated for an additional hour with 150 μg/mL streptomycin to kill not engulfed bacteria. Finally, samples were incubated for 5 min at room temperature with PBS/0.1% Triton-X100 to permeabilize cells and allow the release of internalised bacteria in the supernatants. Bacterial fluorescence was read using a Varioskan Lux microplate reader (Thermo Fisher) as a readout of the uptaken bacteria.

### Live bacterial imaging

ST147_NDM-1_-sfmCherry was grown ON as described before and re-launched until OD_600_ 0.5. The assay was carried out in a final volume of 50 μL containing exponential Kp diluted 1:100, 12.5% BRC and escalating doses of anti-Kp mAbs in PBS. Samples were prepared in 96-well Phenoplate (Perkin Elmer, 6055300) and briefly centrifuged at 1000 rpm to keep bacteria closer to the bottom of the well. Acquisition started immediately after centrifugation and was carried out at 37°C by acquiring frames every 2 min for 2 hours.

### Microscopy and image analysis

Imaging was performed using the Opera Phenix platform (Perkin Elmer) and all samples were acquired using a 63X N.A. 1.15 water objective.

For mAb binding experiments carried out in fixed conditions, 20 fields of view (FOV)/well were selected. For each FOV, five z-stacks separated by a z-step of 0.5 μm were acquired. Imaging was performed in confocal mode by exciting the samples with lasers at 425 nm, 488 nm and 561 nm and collecting the emitted light using bandpass filters 435-480 nm, 500-550 nm and 570-630nm, respectively.

For bacterial live imaging experiments, 3 FOV/well were acquired and three z-stacks separated by a z-step of 0.5 μm were imaged. Imaging was performed in widefield mode by exciting the samples with lasers at 561nm and collecting the light using the bandpass emission filter 570-630 nm.

Images were analyzed with Harmony (v4.9), provided by Perkin Elmer, by using custom-made image analysis pipeline described in Table S5. Individual bacteria were detected using the DAPI channel and filtered according to their morphology based on data available in literature. To measure mAb intensity levels, the mean intensity of the A488 signal was measured within a ROI drawn around each bacterium (**Figure S5**). To measure mAb occupancy area, A488 spots were detected, and their morphology features were measured. A488 spots displaying an area smaller than 0.5 μm^2^ were discarded from the analysis because they were not representative of the distribution of the mAb signals around the bacteria.

### Purification of ST147_NDM-1_ capsule and purification of O-antigen from Kp strains

Capsule and O-antigens were purified according to slightly modified procedures already published ^81,83^.

### Nuclear magnetic resonance spectroscopy (NMR)

10 mg of purified capsule and O antigen material was buffer-exchanged to D_2_O by drying with a SpeedVac and resuspending in 99.9 % D_2_O for 3 times in total. NMR spectra (1H, HSQC, 13C) were recorded at 323 K on Bruker AVANCE NEO and AVANCE MHD NMR spectrometers, operating at 900 and 950 MHz (1H Larmor frequency), respectively, both equipped with triple resonance cryo-probes. The obtained spectra were compared with previously published ones to confirm capsule^82^ and O-antigen^48,84^ identity.

### ELISA on purified Kp O-antigen and capsule

To perform the ELISA, purified O2a-LPS and O2afg -LPS were kindly provided by Chris Whitfield’s group^48^. In addition, in-house purified KL64 capsule and O2a and O1 O-antigens were used to coat high-binding 384-well plates (Greiner ref. 781061) and incubated at 4°C ON. The next day, plates were blocked in PBS-BSA 1% at 37°C for 1h. After blocking, plates were incubated with each mAb as primary antibody at a final concentration of 10 µg/ml for 2h at RT, in the presence of PBS, 1% BSA, and 0.05% Tween-20. Next, anti-human IgG secondary antibody conjugated with Alkaline Phosphatase (Southern Biotech) was added for 1h at 37°C. To detect antigen-mAbs binding, pNPP (p-nitrophenyl phosphate; Sigma-Aldrich) was used as a soluble substrate, and the final reaction was quantified at a wavelength of 405 nm using the Varioskan Lux Reader. After each incubation step, plates were washed three times with 100 µl per well of washing buffer (PBS plus 0.05% Tween-20). Sample buffer and an unrelated mAb were used respectively as blank and negative control, and we considered positive hits those wells showing an OD_405_ value 3-fold higher than the blank. As a positive control, a 1:100 dilution of pooled plasma from different patients was used.

### Immunocompetent ST147_NDM-1_ bacteremia model

This portion of the study was done in the laboratory of Dr. David P. Nicolau and Dr. Kamilia Abdelraouf, at the Center for Anti-infective Research and Development, Hartford Hospital. Specific pathogen-free, female ICR mice weighing 20-22 grams were obtained from Charles River Laboratories, Inc., (Wilmington, MA). The animals were allowed to acclimate for a minimum of 48 h before commencement of experimentation and were provided food and water *ad libitum* ^85,86^. The protocol was reviewed and approved by the Institutional Animal Care and Use Committee at Hartford Hospital. Mice were administered uranyl nitrate 5 mg/kg three days prior to inoculation to produce a controlled degree of renal impairment. ST147_NDM-1_ was previously frozen at −80°C in skim milk (BD BioSciences, Sparks, MD). Prior to mice inoculation, two transfers of the organisms were performed onto Trypticase Soy Agar plates with 5% sheep blood (TSA II™; Becton, Dickinson & Co.; Sparks, MD) and incubated at 37°C for approximately 24 h. After 18-24 h of incubation of the second transfer, a bacterial suspension of approximating 10^6.5^ CFU/ml in 5% hog gastric mucin was made for inoculation. Final inoculum concentrations were confirmed by serial dilution and plating techniques. Septicemia was produced by IP injection of 0.5 ml of the inoculum.

08O09, 05N02 and 05D08 were reconstituted with PBS to the concentrations required to deliver 1, 5, 10, and 20 mg/kg doses based on the mean weight of the study mice population. All dosing solutions were kept on ice while filled syringes with dosing solutions were refrigerated until use. PBS solution with a pH of 7.4 was utilized as the vehicle for dosing control animals throughout the study.

For PRO studies, single dose mAb 08O09, 05N02 or 05D08 of 1, 5, 10, 20 mg/kg were administered through intraperitoneal (IP) route 24 h prior to bacterial inoculation (Figure 5A). For THR studies, single dose 08O09 of 1, 5, 10, 20 mg/kg were administered through intravenous (IV) route 1 h post bacterial inoculation (Figure 5F). For PRO+THR studies, one dose of 1 or 5 mg/kg of 08O09 was administered IP 24 h prior to bacterial inoculation, and a second dose of 08O09 at the same dose was administered IV 1 h post bacterial inoculation (Figure 5H). Each dosing regimen consisted of 10 mice per group. Sham control mice (n=10) received PBS in the same volume, route, and schedule as the study groups. One hour after inoculation, the 0 h control group of 6 mice was euthanized by CO2 exposure followed by cervical dislocation for spleen harvest. Spleens were homogenized and serial dilutions of the samples were plated on agar for CFU determination to confirm the establishment of septicemia. Sham controls as well as the study group mice were followed up for mortality for 96 h. Spleens were harvested and processed as previously described for CFU determination, from the mice that are euthanized or found dead during the observation time. Animals survived till 96 h were euthanized and spleens were also harvested and processed for CFU enumeration.

Efficacy was calculated as the reduction in spleen bacterial load (log_10_ CFU) comparing mice received therapy with sham control. The rate and extent of mortality were recorded and assessed over the four-day (96 h) study period. Bacterial density reductions were analyzed using one-way ANOVA with Tukey’s analyses. Survivals were compared between groups using Kaplan-Meier survival analysis and the log rank test.

Additionally, plasma concentration of 08O09 in uninfected mice were assessed for PRO and THR 5 mg/kg regimens. Mice were prepared as previously described. Plasma was collected through orbital bleeding at baseline for all mice as control samples for analysis. One group of 6 mice were then administered IP single dose of 08O09 5 mg/kg. At 24 h post-dose, plasma was collected through cardiac puncture after euthanasia. Blood samples were collected in vials containing K2 EDTA. The blood was centrifuged and the plasma from each sample was transferred into polypropylene tubes. These tubes were stored at −80 °C until plasma concentrations were determined. The plasma concentrations of 08O09 were determined by quantitative ELISA, as described in previous section. Each plasma concentration was evaluated in 5 replicates and the values from the replicates were averaged for each sample. Values were obtained after 20 min measurement of the plates and interpolated from a sigmoidal curve of 08O09 in presence of mice plasma.

## SUPPLEMENTAL VIDEOS

**SupplementalVideo1_Cluster1 mAb.** Time-lapse imaging of ST147_NDM-1_-sfmCherry treated with 100 µg/mL of anti-Kp mAb 08O09.

**SupplementalVideo2_Cluster2 mAb.** Time-lapse imaging of ST147_NDM-1_-sfmCherry treated with 100 µg/mL of anti-Kp mAb 05N02.

## Notes

### Summary of Updates

The Materials and Methods section has been revised as follows: the paragraphs describing the purification procedures of Klebsiella pneumoniae capsule and O-antigen have been merged into a single one entitled "Purification of ST147NDM-1 capsule and purification of O-antigen from Kp strains".

