## Supplemental information for "Anti-capsule human monoclonal antibodies protect against hypervirulent and pandrug-resistant *Klebsiella pneumoniae*"

**Figure S1. Gating strategy for single cell sorting of total memory B cells and heatmap of ELISA results against ST147$_{NDM-1}$ clinical isolates, related to Figure 1.** (A) Flow cytometry plots reporting gating strategy to identify live cells, lymphocyte population, single cells, CD19$^{+}$ cells (B cells), CD19$^{+}$ CD27$^{+}$ IgD$^{-}$ cells (memory B cells), CD19$^{+}$ CD27$^{+}$ IgD$^{-}$ IgM$^{-}$ B cells (memory B cells expressing IgG, IgA or IgE). (B) ELISA against two different ST147$_{NDM-1}$ clinical isolates (c.i) was performed to confirm mAb binding to Kp surface. Values at least 3-fold times above the OD$_{405}$ blank were considered as positive hits.

A

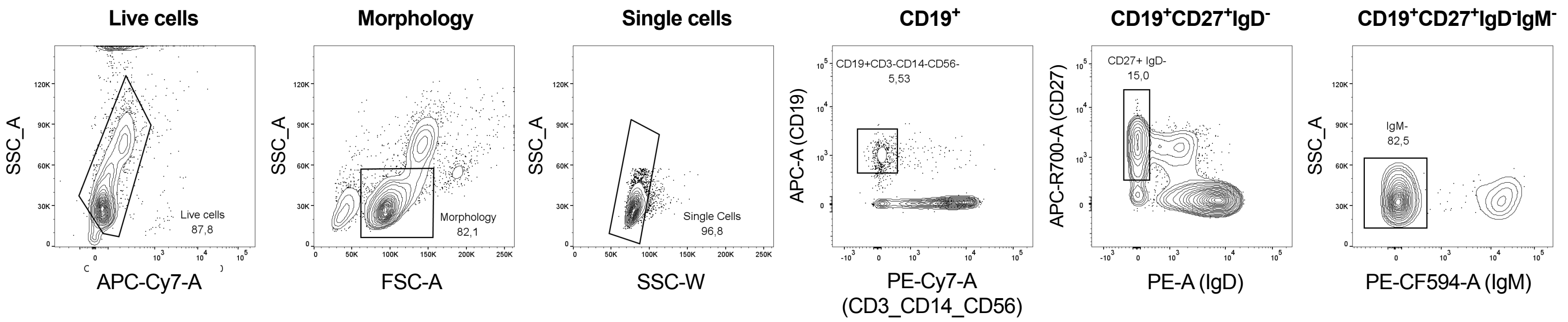


B

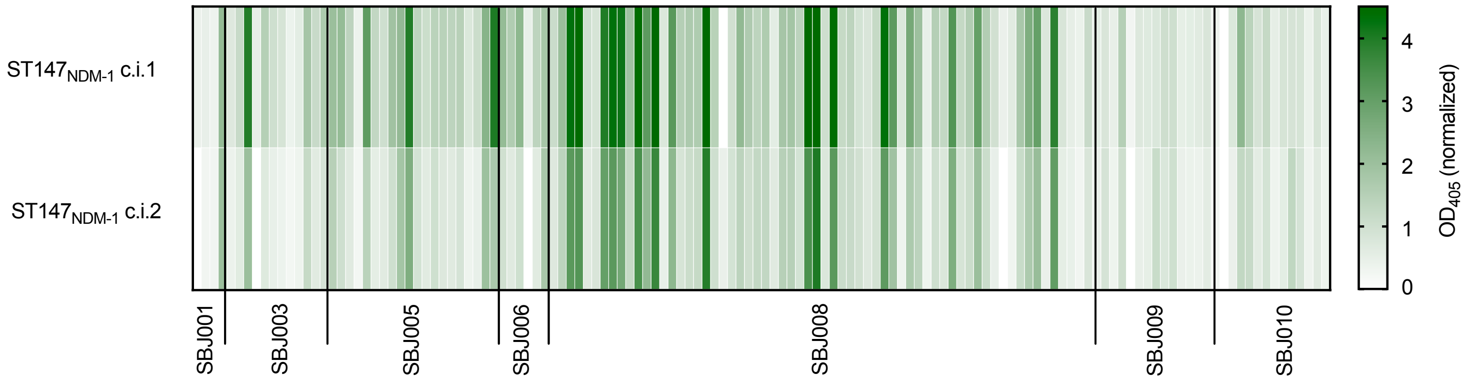


**Supplementary Figure 1**

**Figure S2. Phylogenetic analysis of the Kp strains used for flow cytometry, ELISA and immunoblot experiments and flow cytometry analysis of the mAb binding properties against other Kp species and commensals, related to Figure 2.** (A) Maximum likelihood SNP tree calculated with IQTree, based on the 4189 core genes identified with Roary. K locus, O locus and O type are annotated on the side. Evolutionary model: GTR + F + ASC. (B) Heatmap showing the summary of high-throughput flow cytometry screening of 20 functional anti-Kp mAbs against *Klebsiella oxytoca* and *Klebsiella variicola* isolates, as well as against commensals. MFI values were normalized on no mAb controls and transformed in logarithmic scale.

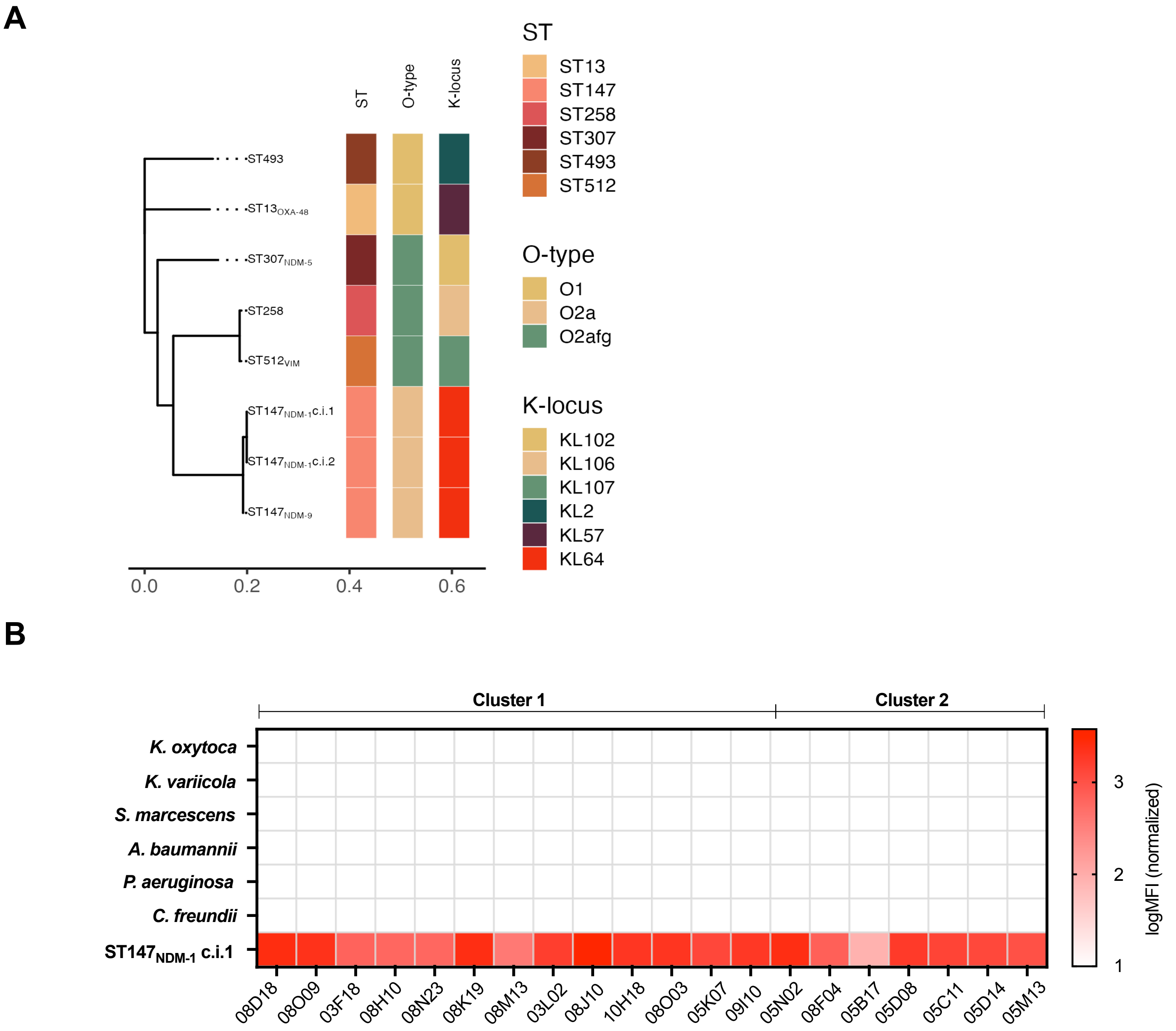


**Supplementary Figure 2**

**Figure S3. Annotation of K and O locus on selected strains, related to Figure 2.** (A) Sequence fragment resulting from the alignment of *wbbO* gene in selected strains, illustrating the single-point deletion at position 474 causing a premature stop codon in $ST147_{NDM-9}$. (B) For each sequence, the presence of K- (blue) and O-locus (red) genes is presented; genes with premature stop codons are visualized as shades.

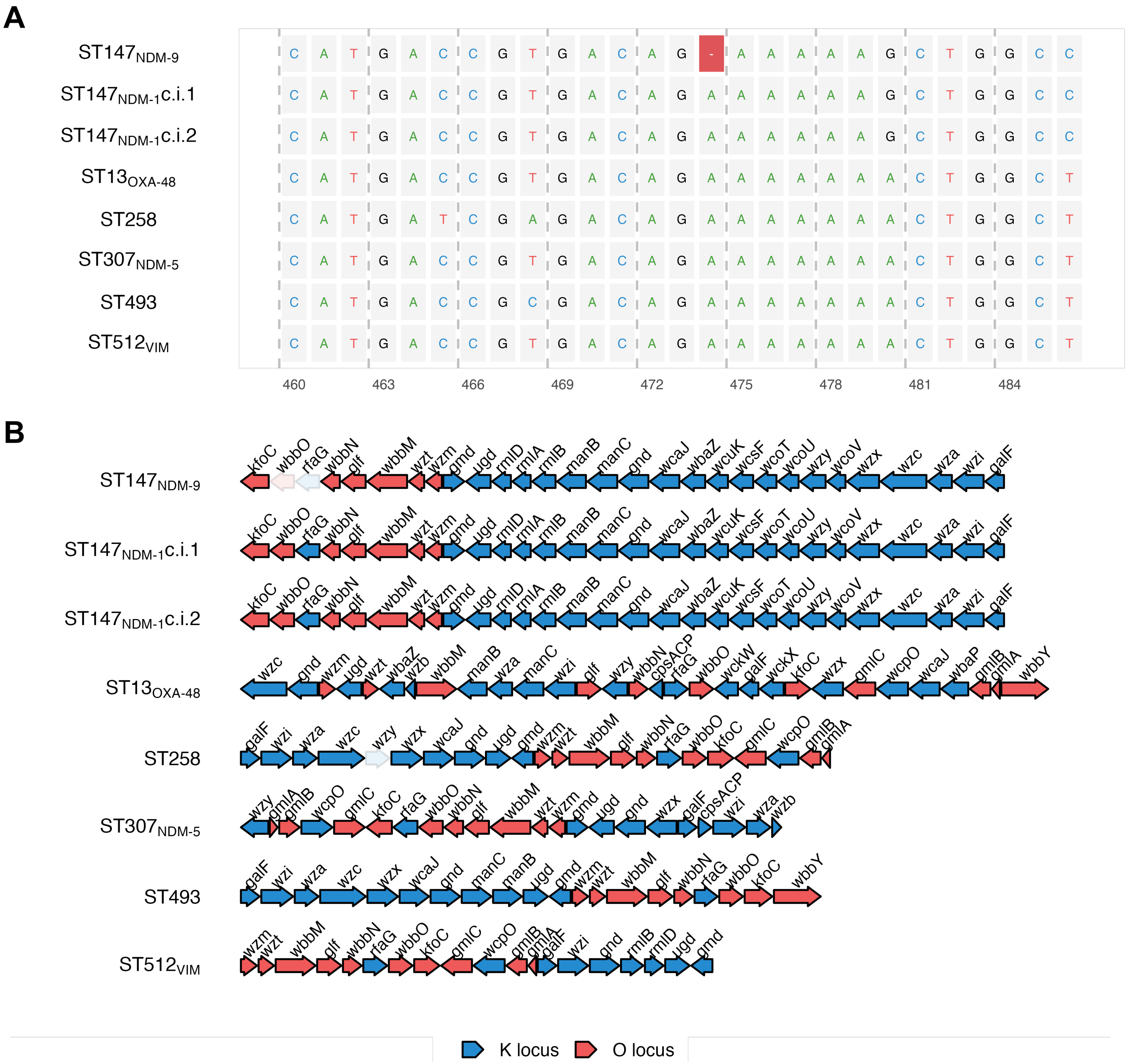


**Supplementary Figure 3**

**Figure S4. Polysaccharide characterization of O-antigen-deficient strain ST147$_{NDM-9}$, purification of capsule type KL64 from ST147$_{NDM-1}$ Kp and purification of different O-Antigen subtypes, related to Figure 2.** (A) SEC-HPLC analysis of total sugar content after hydrolysis with acetic acid. The three peaks correspond to capsule, O-antigen and the core. Three arrows indicate molecular weight standards of 410, 80, and 12 kDa. (B) Silver staining analysis of total sugar extract from ST147$_{NDM-1}$ and ST147$_{NDM-9}$ Kp strains. ST147$_{NDM-9}$ lacks LPS ladder-like signal in the 25-50 kDa range, which is present in the ST147$_{NDM-1}$ loaded sample. (C) SEC-HPLC emission (λ=280nm) and RIU normalized profiles of purified KL64 capsule from ST147$_{NDM-1}$. The three arrows indicate molecular weight standards of 410, 80, and 12 kDa. ST147$_{NDM-1}$ capsular polysaccharide was purified following the protocol reported in Materials and Methods. (D) Theoretical structure of capsule type 64, as described in literature[1] (E) SEC-HPLC profiles of O2a purified from ST147$_{NDM-1}$ and (F) O1v2 purified from ST13$_{OXA-48}$. The three arrows indicate molecular weight standards of 410, 80, and 12 kDa. (G) Molecular structures of O1, O2a, and O2afg O-antigen types as reported in literature[2,3].

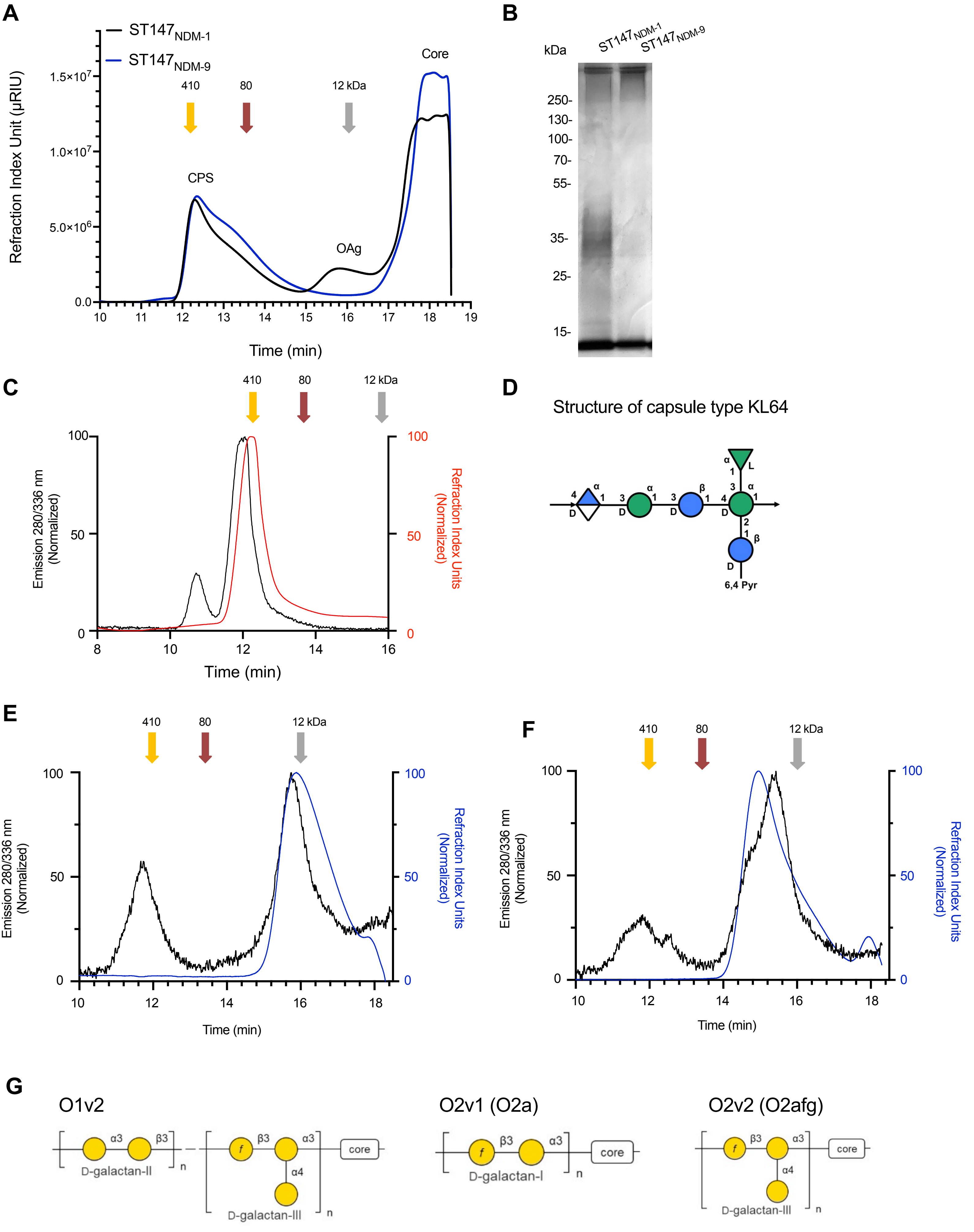


**Supplementary Figure 4**

**Figure S5. ROI definition and spot detection for mAb binding characterization, related to Figure 3.** Images on the left show the binding pattern of A488-labelled anti-Kp mAbs on $ST147_{NDM-1}$ Kp, in the middle the ROI where A488 signal was detected and on the right the morphology of A488 spot. Scale bar 2 µm.

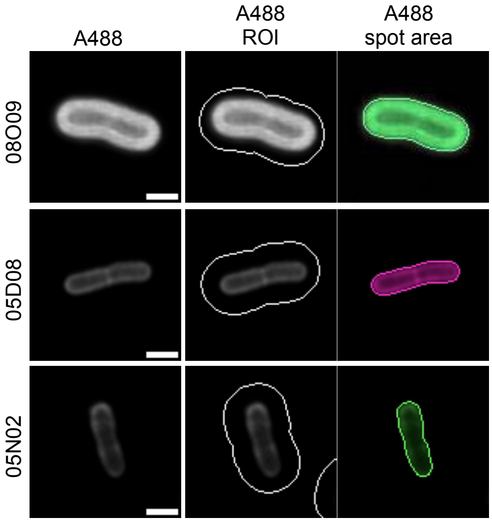


**Supplementary figure 5**

**Figure S6. Sequence analysis of 20 selected mAbs, related to Figure 2**. (A) Frequency of IGHV and IGHJ genes usage (left), IGKV and IGKJ genes usage (right). (B) IGHV and IGHJ genes pairing heatmap (left), IGKV and IGKJ genes pairing heatmap (right). (C-D) Percentage of variable chain identity with respect to the inferred germline. Antibodies similarity was analyzed by calculating a distance matrix using CLUSTAL Omega for the heavy (C) and light (D) chains separately. Dendrogram tips were colored according to the associated mAb cluster.

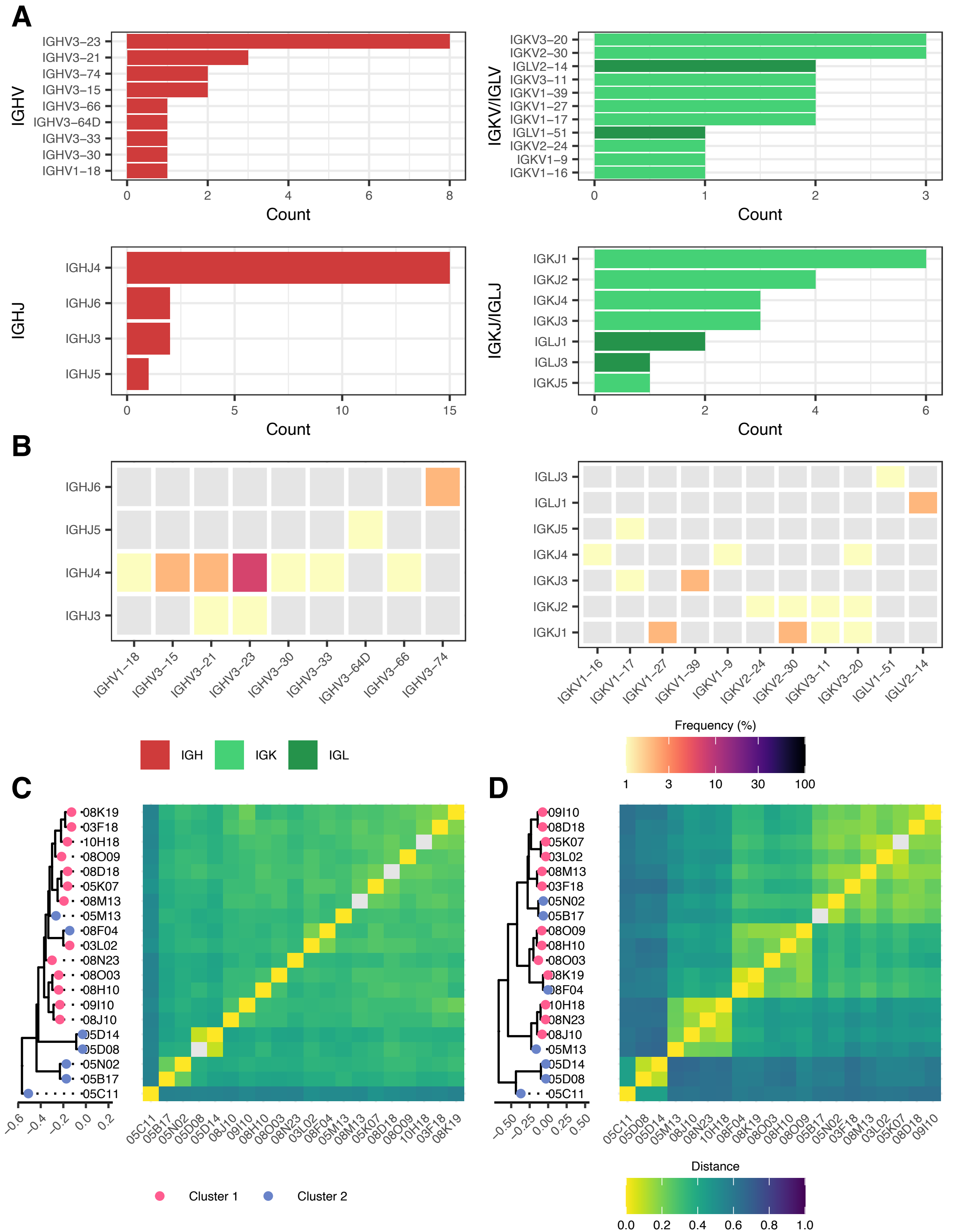
A
IGHV
IGHV3−23
IGHV3−21
IGHV3−74
IGHV3−15
IGHV3−66
IGHV3−64D
IGHV3−33
IGHV3−30
IGHV1−18
Count
IGKV/IGLV
IGKV3−20
IGKV2−30
IGLV2−14
IGKV3−11
IGKV1−39
IGKV1−27
IGKV1−17
IGLV1−51
IGKV2−24
IGKV1−9
IGKV1−16
IGHJ
IGHJ4
IGHJ6
IGHJ3
IGHJ5
IGKJ/IGLJ
IGKJ1
IGKJ2
IGKJ4
IGKJ3
IGLJ1
IGLJ3
IGKJ5
B
IGH
IGK
IGL
Frequency (%)
1
3
10
30
100
C
D
Cluster 1
Cluster 2
Distance
0.0
0.2
0.4
0.6
0.8
1.0
08K19
03F18
10H18
08O09
08D18
05K07
08M13
05M13
08F04
03L02
08N23
08O03
08H10
09I10
08J10
05D14
05D08
05N02
05B17
05C11

**Figure S7. F-SBA profiling and killing efficacy of 20 functional mAbs against complement-sensitive pathogenic Kp strains, related to Figure 4.** F-SBA curves of 20 bactericidal anti-Kp mAbs against $ST147_{NDM-1}$ c.i.1 (A), $ST147_{NDM-1}$ c.i.2 (B), $ST147_{NDM-9}$ (C) and $ST307_{NDM-5}$ (D). Single experiments were normalized to no mAb controls. For each mAb, the bactericidal curve and the associated $IC_{50}$ value were obtained using the [Inhibitor] vs. normalized response, variable slope analysis on GraphPad Prism 10.1.0. $IC_{50}$ values are reported in Figure 4A. (E) The heatmap displays the percentage of reduction in resazurin fluorescence as a readout of bacterial viability, normalized to controls. Results were extrapolated from F-SBA experiments with single mAbs.

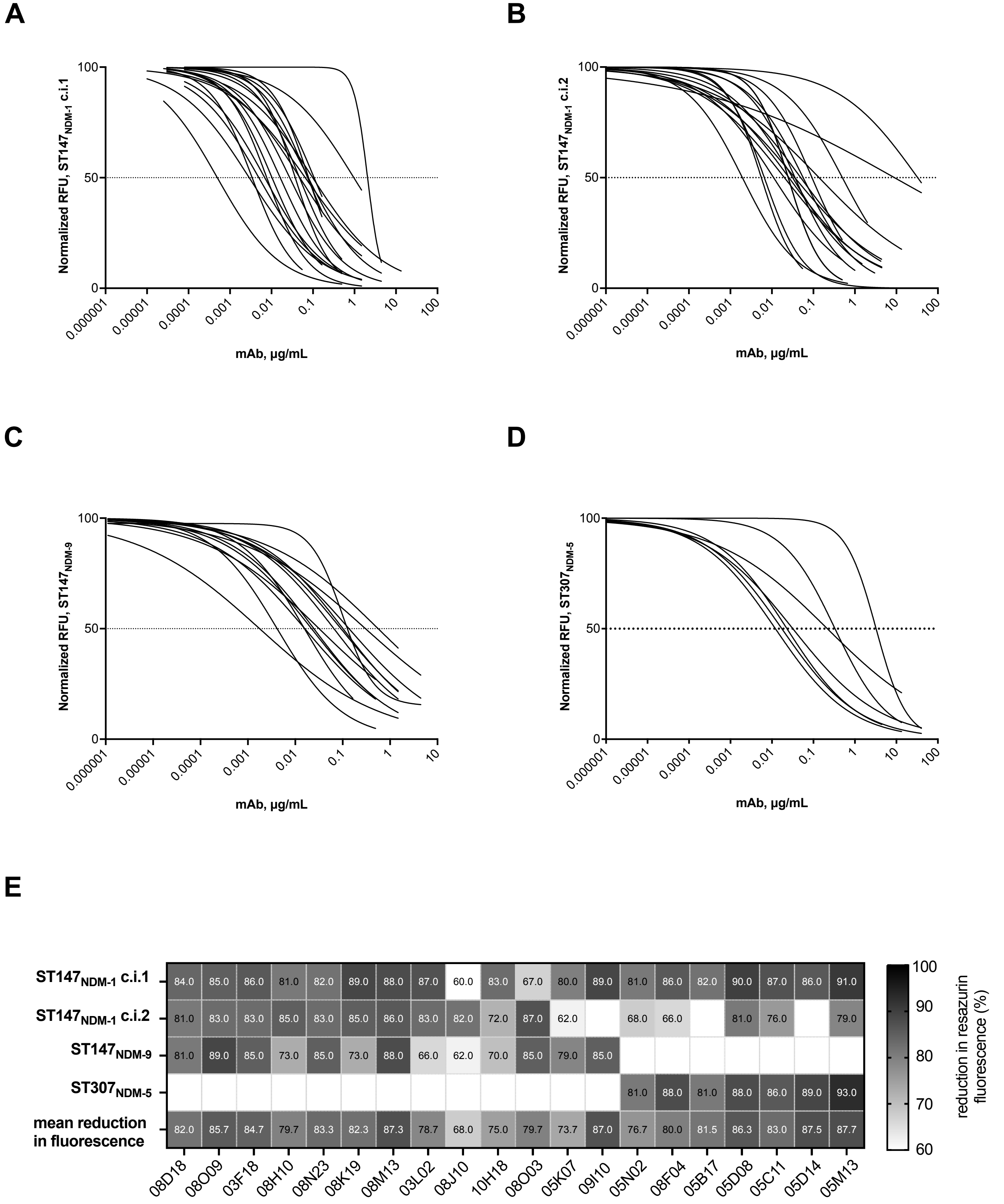


**Supplementary Figure 7**

**Table S1. Patient list.** Information about patients from the Tuscany outbreak enrolled in the study, related to Figure 1

| Patient ID | Age | Date of Bacteremia | Collection Date | Months after infection | Bacterial strain |
|---|---|---|---|---|---|
| **SBJ001** | 67 | 21/08/19 | 21/03/20 | 7 | Klebsiella $ST147_{NDM-1}$ |
| **SBJ003** | 69 | 16/02/20 | 07/10/20 | 8 | Klebsiella $ST147_{NDM-1}$ |
| **SBJ005** | 74 | 02/11/19 | 12/11/20 | 12 | Klebsiella $ST147_{NDM-1}$ |
| **SBJ006** | 77 | 05/11/20 | 04/12/20 | 1 | Klebsiella $ST147_{NDM-1}$ |
| **SBJ008** | 43 | 27/08/20 | 07/01/21 | 5 | Klebsiella $ST147_{NDM-1}$ |
| **SBJ009** | 61 | 11/04/20 | 08/01/21 | 8 | Klebsiella $ST147_{NDM-1}$ |
| **SBJ0010** | 74 | 20/04/20 | 25/01/21 | 8 | Klebsiella $ST147_{NDM-1}$ |

**Table S2. List of primers used in this work, related to Figure 1.**

| Name | Sequence |
|---|---|
| **Heavy chain PCR I primer mix** | |
| L-VH1_VH7 fw | CACTCCCAGGTGCAGCTGGTGCAG |
| L-VH2 fw | TGGGTCTTRTCCCAGGTCACCTTG |
| L-VH 3fw | AAGGTGTCCAGTGTSAGGTGCAG |
| L-VH4_6 fw | GTCCTGTCCCAGGTGCAGCTGCAG |
| L-VH5 fw | GAGTCTGTTCCGAGGTGCAGCTGG |
| IgG CH rev | GTGCCAGGGGGAAGACCGATG |
| IgA CH rev | GCMGAGGCTCAGCGGGAAGAC |
| **Kappa chain PCR I primer mix** | |
| L-VK1 fw | CAGGTGCCAGATGTGHCATCCAG |
| L-VK2 fw | CTGGATCCAGTGSGGATATTGTGATG |
| L-VK3 fw | CCCAGATACCACCGGAGAAATTGTG |
| L-VK4 fw | CTCTGGTGCCTACGGGGACATCGTG |
| L-VK5 fw | CTGATACCAGGGCAGAAACGACAC |
| CK rev 1st | GAACACTCTCCCCTGTTGAAGCTCTTTG |
| **Lambda chain PCR I primer mix** | |
| L-VL1 fw | GGTCCTGGGCCCAGTCTGTGCTG |
| L-VL2 fw | GGTCCTGGGCCCAGTCTGCCCTG |
| L-VL3 fw | TCTGTGRCCTCCTATGAGCTGAC |
| L-VL4_VL5_VL9 fw | CTCTCGCAGCCTGTGCTGACTCA |
| L-VL6 fw | GTTCTTGGGCCAATTTTATGCTG |
| L-VL7 fw | GGTCCAATTCTCAGGCTGTGGTG |
| L-VL8 fw | GAGTGGATTCTCAGACTGTGGTG |
| L-VL10 fw | GTCAGTGGTCCAGGCAGGGCTGAC |
| CL rev 1st | GTGCTCCCTTCATGCGTGACC |
| **Heavy chain PCR II primer mix** | |
| HIFI*_C134_VH1_5_7 | GT ATC ATC CTT TTT CTA GTA GCA ACT GCA ACC GGT GTACATTCC CAG GTG CAG CTG GTG CAG TCT G |
| HIFI*_C134_VH2 | GT ATC ATC CTT TTT CTA GTA GCA ACT GCA ACC GGT GTACATTCC CAG GTC ACC TTG AAG GAG TCT GGT C |
| HIFI*_C134_VH3 | GT ATC ATC CTT TTT CTA GTA GCA ACT GCA ACC GGT GTACATTCC GAG GTG CAG CTG GTG GAG TCT GGG GGA G |
| HIFI*_C134_VH4_6_a | GT ATC ATC CTT TTT CTA GTA GCA ACT GCA ACC GGT GTACATTCC CAG GTG CAG CTG CAG GAG TCG GG |
| HIFI*_C134_VH4_6_b | GT ATC ATC CTT TTT CTA GTA GCA ACT GCA ACC GGT GTACATTCC CAG GTG CAG CTG CAG CAG TGG GG |
| HIFI*_IgG CH rev | CTT GGA GGA GGG TGC CAG GGG GAA GAC CGA TGG GCC CTT GGT GGA RGC |
| HIFI*_IgA CH rev | CTT GGA GGA GGG TGC CAG GGG GAA GAC CGA CTT GGG GCT GGT CGG GGA |
| HIFI*_IgM CH rev | CTT GGA GGA GGG TGC CAG GGG GAA GAC CGA TGG GGC GGA TGC ACT CCC |
| **Kappa chain PCR II primer mix** | |
| HIFI*_C135_VK1 | GT ATC ATC CTT TTT CTA GTA GCA ACT GCA ACC GGT GTACATTCC GCC ATC CAG ATG ACC CAG TCT CCA TC |
| HIFI*_C135_VK2_a | GT ATC ATC CTT TTT CTA GTA GCA ACT GCA ACC GGT GTACATTCC GAT ATT GTG ATG ACC CAG ACT CCA CTC TC |
| HIFI*_C135_VK2_b | GT ATC ATC CTT TTT CTA GTA GCA ACT GCA ACC GGT GTACATTCC GAT ATT GTG ATG ACT CAG TCT CCA CTC TC |
| HIFI*_C135_VK3_a | GT ATC ATC CTT TTT CTA GTA GCA ACT GCA ACC GGT GTACATTCC GAA ATT GTG TTG ACA CAG TCT CCA G |
| HIFI*_C135_VK3_b | GT ATC ATC CTT TTT CTA GTA GCA ACT GCA ACC GGT GTACATTCC GAA ATT GTG ATG ACG CAG TCT CCA G |
| HIFI*_C135_VK4 | GT ATC ATC CTT TTT CTA GTA GCA ACT GCA ACC GGT GTACATTCC GAC ATC GTG ATG ACC CAG TCT CCA G |
| HIFI*_C135_VK5 | GT ATC ATC CTT TTT CTA GTA GCA ACT GCA ACC GGT GTACATTCC GAA ACG ACA CTC ACG CAG TCT CCA G |
| HIFI*_C080_VK_Rev | GA TTT CAA CTG CTC ATC AGA TGG CGG GAA GAT GAA GAC AGA TGG TGC AGC CAC AGT TC |
| **Lambda chain PCR II primer mix** | |
| HIFI*_C080_VL1 | GT ATC ATC CTT TTT CTA GTA GCA ACT GCA ACC GGT TCCTGGGCC CAG TCT GTG CTG ACT CAG CCG CCC TCA G |
| HIFI*_C080_VL2 | GT ATC ATC CTT TTT CTA GTA GCA ACT GCA ACC GGT TCCTGGGCC CAG TCT GCC CTG ACT CAG CCT GCC TCC G |
| HIFI*_C080_VL3_a | GT ATC ATC CTT TTT CTA GTA GCA ACT GCA ACC GGT TCCTGGGCC TCC TAT GAG CTG ACA CAG CCA C |
| HIFI*_C080_VL3_b | GT ATC ATC CTT TTT CTA GTA GCA ACT GCA ACC GGT TCCTGGGCC TCC TAT GAG CTG ACT CAG GAC C |
| HIFI*_C080_VL4 | GT ATC ATC CTT TTT CTA GTA GCA ACT GCA ACC GGT TCCTGGGCC CAG CCT GTG CTG ACT CAA TCG TCC TCT G |
| HIFI*_C080_VL5-9 | GT ATC ATC CTT TTT CTA GTA GCA ACT GCA ACC GGT TCCTGGGCC CAG CCT GTG CTG ACT CAG CCR ACT TC |
| HIFI*_C080_VL6 | GT ATC ATC CTT TTT CTA GTA GCA ACT GCA ACC GGT TCCTGGGCC AAT TTT ATG CTG ACT CAG CCC CAC TC |
| HIFI*_C080_VL7 | GT ATC ATC CTT TTT CTA GTA GCA ACT GCA ACC GGT TCCTGGGCC CAG GCT GTG GTG ACT CAG GAG CCC TC |
| HIFI*_C080_VL8 | GT ATC ATC CTT TTT CTA GTA GCA ACT GCA ACC GGT TCCTGGGCC CAG ACT GTG GTG ACC CAG GAG CCA TC |
| HIFI*_C080_VL10 | GT ATC ATC CTT TTT CTA GTA GCA ACT GCA ACC GGT TCCTGGGCC CAG GCA GGG CTG ACT CAG CCA CCC TCG G |
| HIFI*_CL_generic | G TGT GGC CTT GTT GGC TTG AAG CTC CTC ACT CGA GGG YGG GAA CAG AGT G |
| **Primers to amplify transcriptionally active PCR (TAP) products** | |
| CMV_TAP_FW | TTAGGCACCCCAGGCTTTAC |
| polyA_TAP_Rev | AGATGGTTCTTTCCGCCTCA |

**Table S3 (.xls file). Pangenome presence-absence table of COGs in cluster 1 , related to Figure 2.** The presence/absence table of COGs generated from Roary was annotated with gene ontology (GO), Kegg and PFAM functional annotations, and filtered for COGs present exclusively in strains belonging to the cluster 1.

**Table S4. Sequence analysis of 20 functional mAbs, related to Figure 2**

| Sample | Type | V_usage | D_usage | J_usage | Variable region germline identity, % | CDR3 amino acid length |
|---|---|---|---|---|---|---|
| SBJ03-F18 | light | IGKV1-9*01 | | IGKJ4*01 | 93 706 | 9 |
| SBJ03-F18 | heavy | IGHV3-21*01 | IGHJ3*01 | IGHJ4*02 | 95 222 | 13 |
| SBJ03-L02 | light | IGKV1-27*01 | | IGKJ1*01 | 95 088 | 9 |
| SBJ03-L02 | heavy | IGHV3-30*18 | IGHJ2*01 | IGHJ4*02 | 95 578 | 14 |
| SBJ05-B17 | heavy | IGHV3-74*01 | IGHJ1*01 | IGHJ6*02 | 90 203 | 20 |
| SBJ05-B17 | light | IGKV1-39*01 | | IGKJ3*01 | 89 895 | 9 |
| SBJ05-C11 | heavy | IGHV1-18*01 | IGHJ1*01 | IGHJ4*02 | 89 384 | 13 |
| SBJ05-C11 | light | IGLV1-51*01 | | IGLJ3*02 | 93 515 | 11 |
| SBJ05-D08 | heavy | IGHV3-15*01 | IGHJ5*01 | IGHJ4*02 | 95 286 | 10 |
| SBJ05-D08 | light | IGLV2-14*03 | | IGLJ1*01 | 95 578 | 10 |
| SBJ05-D14 | light | IGLV2-14*01 | | IGLJ1*01 | 93,75 | 10 |
| SBJ05-D14 | heavy | IGHV3-15*01 | IGHJ5*01 | IGHJ4*02 | 92 256 | 10 |
| SBJ05-K07 | heavy | IGHV3-23*04 | IGHJ5*01 | IGHJ4*02 | 96 259 | 18 |
| SBJ05-K07 | light | IGKV1-27*01 | | IGKJ1*01 | 98 592 | 9 |
| SBJ05-M13 | light | IGKV2-24*01 | | IGKJ2*01 | 95 695 | 10 |
| SBJ05-M13 | heavy | IGHV3-64D*06 | IGHJ4*01 | IGHJ5*02 | 90 753 | 6 |
| SBJ05-N02 | heavy | IGHV3-74*01 | IGHJ1*01 | IGHJ6*02 | 92 905 | 20 |
| SBJ05-N02 | light | IGKV1-39*01 | | IGKJ3*01 | 95 819 | 9 |
| SBJ08-D18 | heavy | IGHV3-23*04 | IGHJ5*01 | IGHJ4*02 | 93 493 | 12 |
| SBJ08-D18 | light | IGKV1-17*01 | | IGKJ3*01 | 97 535 | 9 |
| SBJ08-F04 | heavy | IGHV3-33*01 | IGHJ6*01 | IGHJ4*02 | 97 635 | 16 |
| SBJ08-F04 | light | IGKV3-11*01 | | IGKJ1*01 | 99 301 | 9 |
| SBJ08-H10 | heavy | IGHV3-23*04 | IGHJ4*01 | IGHJ4*02 | 91 186 | 12 |
| SBJ08-H10 | light | IGKV3-20*01 | | IGKJ1*01 | 94,81 | 9 |
| SBJ08-J10 | heavy | IGHV3-23*04 | IGHJ1*01 | IGHJ4*02 | 90 785 | 11 |
| SBJ08-J10 | light | IGKV2-30*01 | | IGKJ2*03 | 96 321 | 9 |
| SBJ08-K19 | heavy | IGHV3-21*01 | IGHJ2*01 | IGHJ3*02 | 96 918 | 14 |
| SBJ08-K19 | light | IGKV3-11*01 | | IGKJ2*03 | 99 303 | 10 |
| SBJ08-M13 | heavy | IGHV3-23*04 | IGHJ6*03 | IGHJ4*02 | 93 836 | 15 |
| SBJ08-M13 | light | IGKV1-16*02 | | IGKJ4*01 | 98 592 | 9 |
| SBJ08-N23 | heavy | IGHV3-66*01 | IGHJ6*01 | IGHJ4*02 | 92,15 | 14 |
| SBJ08-N23 | light | IGKV2-30*02 | | IGKJ1*01 | 96 296 | 8 |
| SBJ08-O03 | heavy | IGHV3-23*04 | IGHJ3*01 | IGHJ4*02 | 93 878 | 13 |
| SBJ08-O03 | light | IGKV3-20*01 | | IGKJ4*01 | 97 544 | 12 |
| SBJ08-O09 | light | IGKV3-20*01 | | IGKJ2*03 | 96 194 | 10 |
| SBJ08-O09 | heavy | IGHV3-23*04 | IGHJ2*01 | IGHJ3*01 | 94 218 | 12 |
| SBJ09-I10 | heavy | IGHV3-23*04 | IGHJ1*01 | IGHJ4*02 | 89 492 | 14 |
| SBJ09-I10 | light | IGKV1-17*01 | | IGKJ5*01 | 97 183 | 9 |
| SBJ10-H18 | heavy | IGHV3-21*01 | IGHJ6*01 | IGHJ4*02 | 95 254 | 12 |
| SBJ10-H18 | light | IGKV2-30*01 | | IGKJ1*01 | 97 674 | 9 |

**Table S5 (.xls file). Image analysis pipeline developed with Harmony 4.9 for mAb binding characterization, related to Figure 3.**
